# Culturable approach to rice-root associated bacteria in Burkina Faso: diversity, plant growth-promoting rhizobacteria properties and cross-comparison with metabarcoding data

**DOI:** 10.1101/2023.05.30.542993

**Authors:** Moussa Sondo, Issa Wonni, Kadidia Koïta, Isabelle Rimbault, Mariam Barro, Charlotte Tollenaere, Lionel Moulin, Agnieszka Klonowska

## Abstract

Plant-associated bacteria are essential partners in plant health and development. In addition to taking advantage of the rapid advances recently achieved in high-throughput sequencing approaches, studies on plant-microbiome interactions require experiments with culturable bacteria. A study on the rice root microbiome was recently initiated in Burkina Faso. As a follow up, the aim of the present study was to develop a collection of corresponding rice root-associated bacteria covering maximum diversity so as, to be able to assess the diversity of the collection based on the culture medium used, and to describe the taxonomy, phenotype and abundance of selected isolates in the rice microbiome. More than 3,000 isolates were obtained using five culture media (TSA, NGN, NFb, PCAT, Baz). The 16S rRNA fragment sequencing of 1,013 selected working collection isolates showed that our working collection covered four bacterial phyla (Proteobacteria, Firmicutes, Actinobacteria and Bacteroidetes) and represented 33% of the previously described diversity of the rice root microbiome at the order level. Phenotypic *in vitro* analysis of the plant growth promoting capacity of the isolates revealed an overall ammonium production and auxin biosynthesis capacity, while siderophore production and phosphate solubilisation were enriched in *Burkholderia*, *Ralstonia*, *Acinetobacter* and *Pseudomonas* species. Of 45 representative isolates screened for growth promotion on seedlings of two rice cultivars, five showed an ability to improve the growth of both cultivars, while five others were effective on only one cultivar. The best results were obtained with *Pseudomonas taiwanensis* ABIP 2315 and *Azorhizobium caulinodans* ABIP 1219, which increased seedling growth by 158% and 47%, respectively. Among the 14 best performing isolates, eight appeared to be abundant in the rice root microbiome dataset from previous study. The findings of this research contribute to the functional description of rice root-associated bacteria and their potential importance for plants by providing, for the first time, insight into their prevalence in the rice root microbiome.

## INTRODUCTION

Plant health and development are highly influenced by the associated microbiomes. The conventional reductionist approach based on dual plant-microorganism interaction analyses has showcased the contributions of many microorganisms in improving plant growth. These involve enhancment of nutrient pools, nutrient-use efficiency, phytohormone production, adaptation to changing environments, or plant protection from infection by inducing systemic resistance or antagonistic effects on phytopathogens (1–3). With the improvement and cost reduction of high-throughput sequencing (HTS) technologies, new holistic approaches using amplicon barcoding of taxonomic markers or metagenomics have been developed to study the diversity of plant-associated microbiota and their role and function in plant health (3). In the last decade, plant microbiomes have been extensively studied, thereby revealing their unknown diversity in model plants (4,5) and crops (6–9). These studies have revealed the main effects of soil types, cultural practices, plant biodiversity, plant developmental state and climatic conditions on the associated microbial population diversity and structure. Several studies have highlighted the importance of taxonomic clades whose abundance shifts in response to environmental factors (10), but also the putative role of certain taxonomic units belonging to core or specific microbiota (11) or considered as hubs in co-occurrence networks (12). These approaches have also revealed microbiota-mediated disease resistance in plants (reviewed in (13). To further investigate the role of selected taxa (as a clade or unit), functional modelling based on 16S amplicons allows predictive analysis of the population functional potential (14), while metagenomics provides information on gene content (15–17). However, these are only extrapolative analyses that obviously cannot predict differences in microbial function. There is hence still a gap in our knowledge on the activity, function and interaction of the microbiota with host plants, on the function of individual taxa within the population and, finally, on interactions between the microorganisms themselves. Addressing these questions represents a new challenge to go beyond the predictive understanding of gene function, via the metagenome, by developing a metaphenome that describes the functions performed by the microbiome and its physiological, metabolic and interactive status (18–20). In this regard, the use of synthetic communities (SynCom) is a trade-off between reductionist and holistic approaches aimed at unravelling the interactions between the microbiome and host plants on a large scale (21–23). Yet this approach requires numerous extensive collections of microorganisms associated with different plant compartments (rhizosphere, rhizoplan, endosphere, phyllosphere) and knowledge on their relative functions. An ideal SynCom would be derived from each specific plant and its associated environment, while avoiding mixtures of microbes from different origins, as microbial taxa have adaptations to local conditions and specific interactions with host species or other microbiome components (22). Ding *et al.* (24) have suggested that the rice rhizosphere microbiome is markedly different (at the phylum level) from that of other plants (maize, soybean, potato, *Populus* and *Arabidopsis*), partly because growing rice under oxic-anoxic conditions (due to flooding) favours anaerobic microbe development. Differences have been observed between bacterial (higher proportions of Deltaproteobacteria and Chloroflexi), archaeal (higher proportions of Euryarchaeota) and fungal (higher proportions of Chytridiomycota and Zygomycota) populations (24). Efforts to isolate rice-associated bacteria with the aim of describing isolates with plant growth promotion (PGP) features (including free nitrogen fixation) and/or biocontrol capabilities began three decades ago (25–37). However, to the best of our knowledge, only two large bacterial collections on rice have been recently described, i.e. one in Europe (Italy, (38)) and another in Asia (China, (39)). Otherwise, many studies have been devoted to studying specific rice-associated prokaryotes such as methanogenic archaea and methanotrophic bacteria (40,41) and bacteria whose activity causes iron toxicity in lowland areas (42) and others which can mitigate the effects of this toxicity (43,44).

The isolation of plant-associated bacteria has long been based mainly on their growth capacity on common culture media, followed by selection of isolates for plant growth promotion and/or biocontrol capabilities—this has resulted in collections with low taxonomic or functional diversity (45). Community-based cultures (CBC) (46) and high-throughput bacterial cultivation methods (39,47) have facilitated larger-scale capture and manipulation of the bacterial community. In this regard, maximising bacterial diversity has been enhanced by adapting different culture media to specific soil and plant environmental conditions (reviewed in (45) and in (48)). However, these are step by step improvements which had not yet been used in synergy. For plant associated microbiota, the overall recovery of bacterial diversity could therefore range from 4.6% for rice endophytes recovered by (38), compared with barcoding data at the genus level, to 35.3% for culturable rice root-associated bacteria reported by (49), compared with barcoding data at the family level. This proportion was as high as 65% for *Arabidopsis thaliana* root-derived microbiota culture collection (29) and 71.7% for rice root bacterial collection (39) when rare species were removed from the barcoding data (≥0.1% of the relative abundance).

Rice is a staple food for more than half of the world’s population, a vast majority of whom live in developing countries. In Africa, rice accounts for 25% of overall cereal consumption, second only to maize (50). Over the last 20 years, there has been a major surge in rice consumption especially in West Africa as a consequence of demographic growth and habit changes due to urbanisation (preference for fast-prepared food such as rice, compared to other cereals such as millet or sorghum). In reaction to the 2008 food price crisis, West African states increased local rice production with the aim of decreasing their dependency on the worldwide rice market (51). In Burkina Faso, in particular, rice-growing areas increased four-fold between 2006 and 2021 (2023, FAO: fao.org/faostat/). This increase was associated with rice production intensification which, alongside the increased cropping area led to major agricultural changes. In the light of the importance of rice and its specific cultivation conditions that substantially influence its microbiota (24) in the sustainable agriculture context, major efforts are needed to describe and study rice-associated microorganisms, especially in Africa where few such studies have been carried out. While several studies have focused on rice breeding (52), cultivated rice diversity (53) and rice phytopathogens (54–57), studies have only recently been conducted to analyse the rice microbiome (37,49,58). Barro *et al.* (58) showed that the rice production system is a major driver of the microbiome structure in rice fields in Burkina Faso. Higher rhizosphere prokaryotic community diversity and more complex co-occurrence networks have been found in irrigated systems compared to rainfed lowlands, while the opposite pattern has been noted for fungal communities (higher richness in rainfed lowlands). (49) analysed bias related to bacteria cultivability and showed that the use of three popular bacterial media (TSA, NGN; NFb) enabled the recovery of about 1/3 of the total microbiota.

In this study, we aimed to develop a large collection of culturable rice-associated bacteria from different rice fields in Burkina Faso to serve as a valuable genetic resource for screening biostimulant and biocontrol strains. Over 3,800 rice root-associated bacteria were isolated from five bacterial culture media. A working collection of 1,013 isolates was characterized at the taxonomic (by 16S fragment sequencing) and phenotypic (*in vitro* PGP capacity) levels and cross-referenced with rice root microbiome barcode data (16S amplicons) obtained from the same rice fields (58) so as to identify core and hub taxa in the collection. A set of 45 representative strains of the culturable diversity was screened in controlled conditions for plant growth promotion capacities on two rice cultivars popular in Burkina Faso. The best performing isolates were then analysed for their relative abundance in the rice root microbiome dataset.

## MATERIAL AND METHODS

### Rice sampling

*Oryza sativa* ssp. roots were collected from 22 fields in two regions of western Burkina Faso, i.e. Hauts-Bassins (Houet provinces, Bama (irrigated rice, IR) and Badala (rainfed lowland rice, RL) cropping areas) and Cascades (Comoe provinces, Karfiguela (IR) and Tengrela (RL) cropping areas), as shown in Fig 1. Half of these fields have already been described by Barro et al. with regard to agricultural practices, rice diseases, rice genetic diversity and root-associated microbiota (56,59,60). Rice sampling was authorized by a national agreement between the Burkina Faso government and farmers within the framework of a rice productivity improvement program involving INERA. In addition, an “Agreement on the transfer of biological material” was signed between INERA and IRD concerning the samples transfer for this study. Rice roots were sampled over three years, from 2017 to 2019, at different rice development stages (seedling, maximum tillering and flowering-heading), from June to December, to maximise the bacterial isolate collection diversity (S1 Table). Per field, three rice root systems 10 m apart were excavated (20 cm deep), shaken by hand to remove non-adherent soil, placed together in a plastic bag, transported to the laboratory and stored at 10°C until processing.

**Fig 1.**
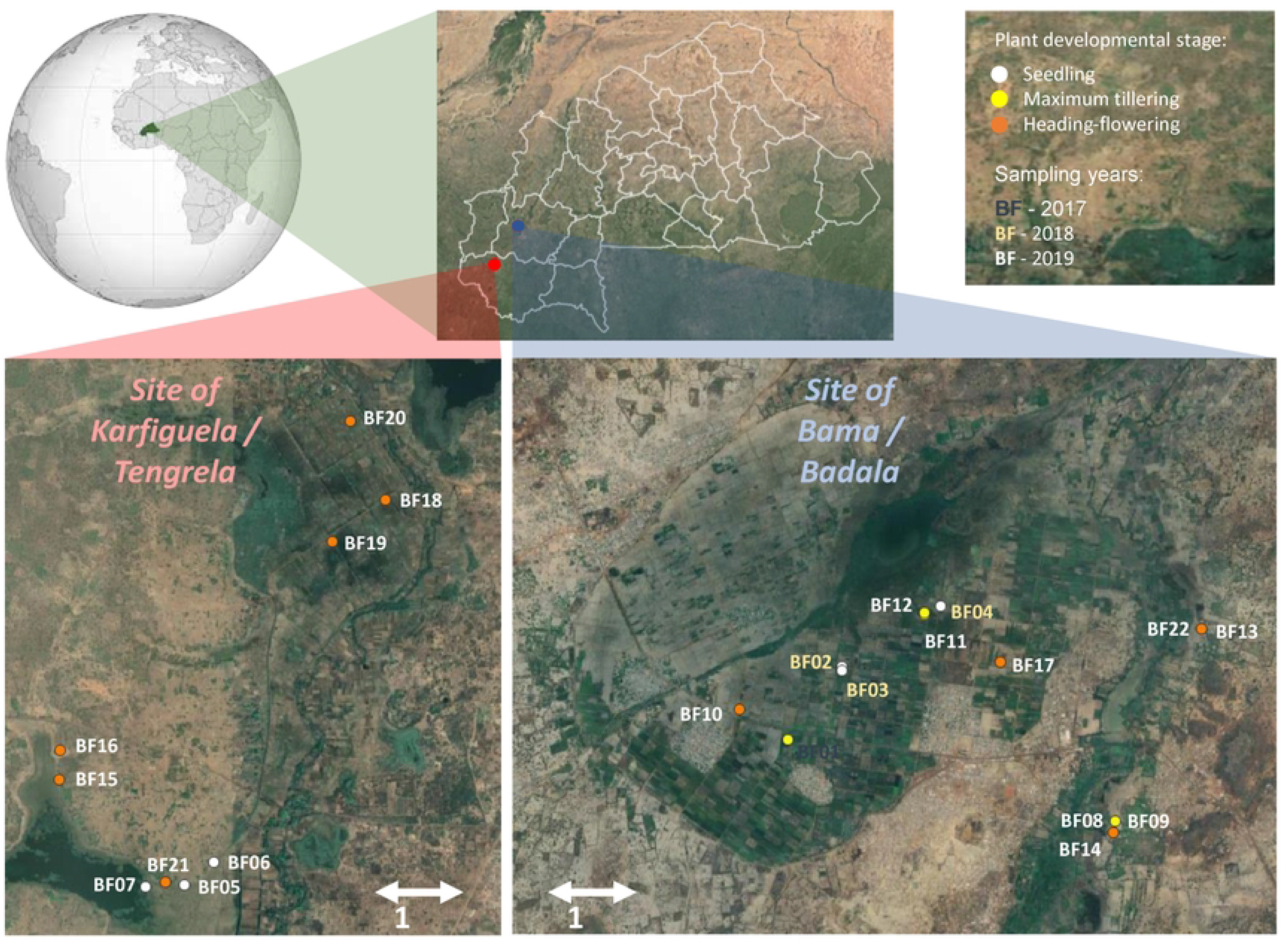
Location of sampled fields within the Haut-Bassins and Cascades regions of western Burkina Faso (sampled in 2017, 2018 and 2019) Table 1 provides a detailed description of the fields.

**Table 1.**
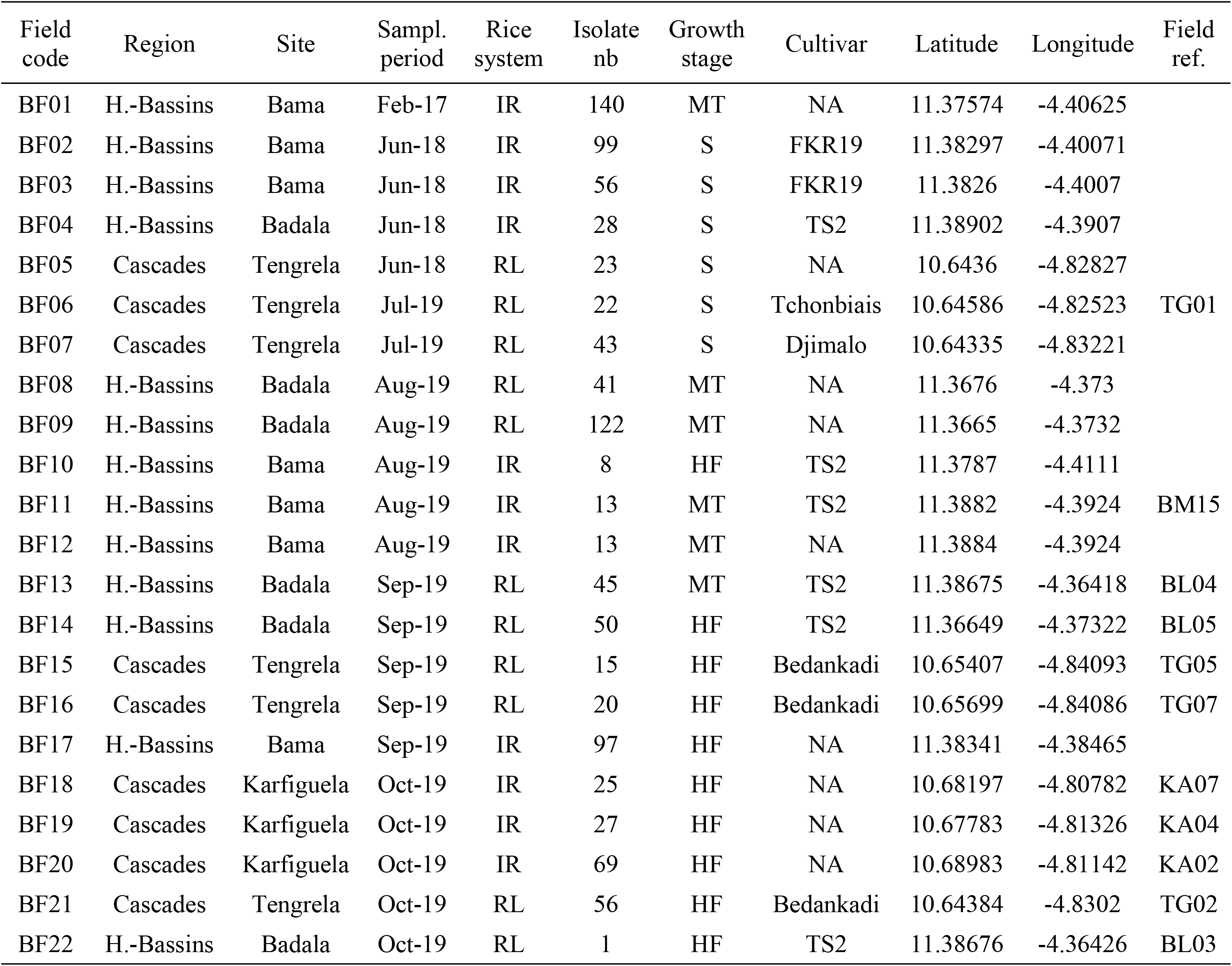
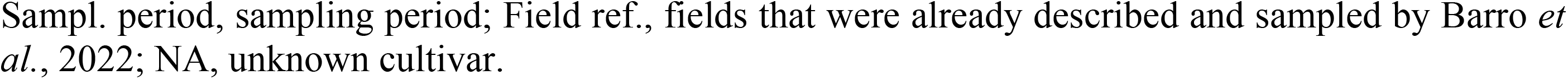
Geographic and diachronic description of sampled fields and rice developmental stages.

| Field code | Region | Site | Sampl. period | Rice system | Isolate nb | Growth stage | Cultivar | Latitude | Longitude | Field ref. |
| --- | --- | --- | --- | --- | --- | --- | --- | --- | --- | --- |
| BF01 | H.-Bassins | Bama | Feb-17 | IR | 140 | MT | NA | 11.37574 | -4.40625 |  |
| BF02 | H.-Bassins | Bama | Jun-18 | IR | 99 | S | FKR19 | 11.38297 | -4.40071 |  |
| BF03 | H.-Bassins | Bama | Jun-18 | IR | 56 | S | FKR19 | 11.3826 | -4.4007 |  |
| BF04 | H.-Bassins | Badala | Jun-18 | IR | 28 | S | TS2 | 11.38902 | -4.3907 |  |
| BF05 | Cascades | Tengrela | Jun-18 | RL | 23 | S | NA | 10.6436 | -4.82827 |  |
| BF06 | Cascades | Tengrela | Jul-19 | RL | 22 | S | Tchonbiais | 10.64586 | -4.82523 | TG01 |
| BF07 | Cascades | Tengrela | Jul-19 | RL | 43 | S | Djimalo | 10.64335 | -4.83221 |  |
| BF08 | H.-Bassins | Badala | Aug-19 | RL | 41 | MT | NA | 11.3676 | -4.373 |  |
| BF09 | H.-Bassins | Badala | Aug-19 | RL | 122 | MT | NA | 11.3665 | -4.3732 |  |
| BF10 | H.-Bassins | Bama | Aug-19 | IR | 8 | HF | TS2 | 11.3787 | -4.4111 |  |
| BF11 | H.-Bassins | Bama | Aug-19 | IR | 13 | MT | TS2 | 11.3882 | -4.3924 | BM15 |
| BF12 | H.-Bassins | Bama | Aug-19 | IR | 13 | MT | NA | 11.3884 | -4.3924 |  |
| BF13 | H.-Bassins | Badala | Sep-19 | RL | 45 | MT | TS2 | 11.38675 | -4.36418 | BL04 |
| BF14 | H.-Bassins | Badala | Sep-19 | RL | 50 | HF | TS2 | 11.36649 | -4.37322 | BL05 |
| BF15 | Cascades | Tengrela | Sep-19 | RL | 15 | HF | Bedankadi | 10.65407 | -4.84093 | TG05 |
| BF16 | Cascades | Tengrela | Sep-19 | RL | 20 | HF | Bedankadi | 10.65699 | -4.84086 | TG07 |
| BF17 | H.-Bassins | Bama | Sep-19 | IR | 97 | HF | NA | 11.38341 | -4.38465 |  |
| BF18 | Cascades | Karfiguela | Oct-19 | IR | 25 | HF | NA | 10.68197 | -4.80782 | KA07 |
| BF19 | Cascades | Karfiguela | Oct-19 | IR | 27 | HF | NA | 10.67783 | -4.81326 | KA04 |
| BF20 | Cascades | Karfiguela | Oct-19 | IR | 69 | HF | NA | 10.68983 | -4.81142 | KA02 |
| BF21 | Cascades | Tengrela | Oct-19 | RL | 56 | HF | Bedankadi | 10.64384 | -4.8302 | TG02 |
| BF22 | H.-Bassins | Badala | Oct-19 | RL | 1 | HF | TS2 | 11.38676 | -4.36426 | BL03 |

Sampl. period, sampling period; Field ref., fields that were already described and sampled by Barro *et al.*, 2022; NA, unknown cultivar.

Four to six days after sampling, roots were rinsed with tap water, placed in a 50 ml Falcon tube containing 30 ml sterile MilliQ water and vortexed for 5 min to separate roots from rhizosphere soil and rinsed three times. Nodal, basal and primordial roots were cut into 1 cm fragments and four different fragments were placed in two separate 2 ml Eppendorf tubes for isolation of: a) total root associated bacteria, and b) root endophytic bacteria. A pre-treatment step was carried out for endophytic bacteria. Roots were surface sterilised by soaking in 4.5% sodium hypochlorite for 1 min and then transferred to sterile water for 1 min. Root fragments from the total and endophytic bacteria treatments were then rinsed four times with sterile water, placed in corresponding 2 ml screw-capped tubes containing 1 mL of sterile water and a 6.3 mm ceramic sterile bead (MP Biomedicals) and homogenised in a FastPrep (MP Biomedicals) at 240 rpm for 2 x 40 s. The series of 10^-2^ to 10^-5^ dilutions was performed with the ground root suspensions. Fifty µL of each dilution were spread on solid culture medium plates and only the 10^-1^ dilution was inoculated in semi-solid medium tubes.

Five culture media with different carbon and nitrogen sources were used to maximise the isolated bacteria diversity. We used a universal non-selective Tryptone Soy Agar (TSA, Sigma) medium at 10% (TSA10) and 50% (TSA50) and four nitrogen-free media for the isolation of potential nitrogen fixers, i.e. Norris Glucose Nitrogen Free Medium (NGN) (61), modified PCAT without nitrogen, (62), and two semi-solid media, including NFb (63,64) and BAz (65). The compositions of the above culture media were as follows: TSA (g/L), tryptone, 1.5; soya peptone, 0.5; NaCl, 1.5; pH adjusted to 7.3; NGN (g/L), K_2_HPO_4_, 1.0; CaCO_3_, 1.0; NaCl, 0.2; MgSO_4_·7H_2_O, 0.2; FeSO_4_·7H_2_O, 0.01; Na_2_MoO_4_.2H_2_O, 0.005; 10 g glucose as carbon source, and pH 7; PCAT (g/L), MgSO_4_, 0.1; azelaic acid, 2.0 (sole carbon source); K_2_HPO_4_, 4.0; KH_2_PO_4_, 4.0; yeast extract, 0.02; pH adjusted to 5.7; NFb (g/L), K_2_HPO_4_, 0.5; MgSO4.7H_2_O, 0.2; NaCl, 0.1; CaCl_2_. 2H_2_O, 0.02; KOH, 4.5; malic acid, 5.0; 2 mL micronutrient solution ((g/L) CuSO_4_.5H_2_O, 0.04; ZnSO_4_.7H_2_O, 0.12; H_3_BO_3_, 1.4; Na_2_MoO_4_.2H_2_O, 1.0; MnSO_4_.H_2_O, 1.175; 2 mL bromothymol blue (5 g/L in 0.2 N KOH), 4 mL FeEDTA (solution 16.4 g/L), 1 mL vitamin solution ((mg/0.1 L) 10 mg biotin, 20 mg pyridoxal-HCl) with pH adjusted to 6.5; BAz (in g/L): azelaic acid, 2.0; K_2_HPO_4_, 0.4; KH_2_PO_4_, 0.4; MgSO_4_.7H_2_O, 0.2; CaCl_2_, 0.02; Na_2_MoO_4_.H2O, 0.002; FeCl_3_, 0.01; bromothymol blue, 0.075; and pH adjusted with KOH to 5.7. All solid media were amended with cycloheximide (200 mg/l) as antifungal agent. Solid and the semi-solid media were obtained by adding 2% and 0.16% of agar, respectively. Only plates containing 20 to 300 CFU were retained for bacterial isolation. Bacterial colonies with different shapes, sizes and colours were purified separately by subculture on TSA50 petri dishes or, in the case of semi-solid NFb and Baz, on solid NFb and BAz media. The pure isolates were then cultured in 4 mL tryptone soy broth 50% (TSB50) for 12 to 24 h at 28°C with shaking. Two millilitres were used for long-term storage of the isolates at −80°C (in 20% glycerol) in individual cryotubes and in 96-well microplates, and the remaining 2 mL of bacterial culture was used for taxonomic characterization of the isolates.

### Taxonomic characterization of isolates

Total bacterial DNA was obtained from isolates using a rapid proteinase K method, as described by (66). The 16S rRNA fragment was amplified using 0.625 units of GoTaq DNA Polymerase (Promega) according to the manufacturer’s instructions, 20 pmol of each forward primer (FGPS1509: AAGGAGGGGATCCAGCCGCA) and reverse primer (FGPS6: GGAGAGTTAGATCTTGGCTCAG) (Normand et al. 1992), as previously described in (Moulin et al. 2001). A routine PCR protocol was used for DNA amplification based on initial denaturation at 94°C for 2 min, then 35 cycles of 30 s denaturation at 94°C, 30 s hybridisation at 55°C steps and a 1 min final extension step at 72°C. An almost full-length 16S rRNA gene was amplified with primers and sequenced with an internal primer (16S-1080.r: GGGACTTAACCCAACATCT). The obtained sequences were assigned at the taxonomic level using BLAST using R software and then deposited in GenBank (https://ncbi.nlm.nih.gov/genbank/) under accession numbers OQ061502 to OQ062366 (see details in Supplementary S1 Table).

### Functional characterization of isolates

Isolates were screened for the following plant growth promotion traits: indole-3-acetic acid (IAA) synthesis, siderophore production, solubilisation of an insoluble phosphate source (Ca_3_(PO_4_)_2_) and ammonium biosynthesis. We screened the large collection of isolates (1,013 clones) by adapting previously published protocols for testing in 96-well culture plates. Isolates were grown in 96-well microplates containing 150 μL TSB50 for 24 h at 28°C at 160 rpm on an orbital shaker. When 0.9 OD was reached, bacterial isolates were either plated on solid medium (square plates) using a 96-pin microplate replicator (Boekel Scientific) for siderophore and phosphate solubilisation assays, or inoculated in 96-well microplates containing 150 μL TSB50 for IAA and ammonia production assays.

IAA production was determined by the Salkowski colourimetric assay, as described in (67). The 24 h bacterial culture, which was performed in 96-well microplates containing 150 µL TSA50, was supplemented with 5 mM liquid L-tryptophan and 30 µL Salkowski reagent (68). Auxin production was detected after 1 h incubation (69). The ability to solubilise inorganic phosphate was assessed on solid TriCalcium orthoPhosphate (TCP) medium (70) supplemented with Ca_3_(PO_4_)_2_ and either KNO_3_ or NH_4_Cl. Isolates were considered positive if they were able to solubilise phosphate in both media after 7 days incubation at 28°C. The TCP medium composition was as follows (g/L): 10 g glucose, 4 g Ca_3_(PO_4_)_2_, 1 g NaCl, 10 g KNO_3_ or 5 g NH_4_Cl, 1 g MgSO_4_.7H_2_O, 20 g agar and pH adjusted to 7.2. Siderophore production was determined on solid yeast manitol agar (YMA) medium supplemented with 10% (v/v) CAS solution; the development of a yellow or orange halo around the bacterial colonies indicated the presence of siderophores. For ammonia production, 100 μL of isolates grown in TSA50 liquid medium were subcultured in peptone water (71) and incubated at 28°C for 48 h, and then 10 μL of Nessler’s reagent (Sigma-Aldrich) was added and ammonium production was detected 20 min later (72). All tests were performed in triplicate for each strain.

### Greenhouse trial

Seedling trays with eight cavities (22 cm^3^) each were filled with potting soil and tamped. Two microplates per condition were placed in each tray and the whole set was sprinkled with osmosis water. Seeds of two rice cultivars from Burkina Faso (FKR64 (TS2) and FKR84 (Orylux 6)) were soaked in tap water for 24 h. Two well watered seeds were transplanted in the middle of each cell. Seedlings were removed at germination to ensure one plant per cavity and homogeneity of plants. The greenhouse experiment was conducted on two large trays, each with its own control, under alternating conditions of 12 h light (day) at 28°C and 80% humidity and 12 h darkness (night) at 28°C and 70% humidity.

For bacterial inoculum preparation, 45 strains were removed from the freezer (at −80°C) and inoculated on TSA50 solid medium. After 48 h incubation at 28°C, bacterial colonies were subcultured 16 hrs at 28°C in 20 mL of TSB liquid medium in a 50 mL Falcon tube. The bacterial culture was centrifuged at 4,000 rpm for 10 min, the supernatant discarded and the pellet resuspended in sterile osmosis water. The bacterial suspension optical densities (OD600) were then adjusted to a final OD of 0.5. To assess the plant growth promotion effect, 1 mL of prepared bacterial suspension was pipetted into the soil at the root base of 5-day-old seedlings and whole plants were collected 10 days later to record the plant size (Size), root dry weight (RW) and leaf dry weight (LW). Each tray, i.e. a 16-well dish, corresponded to a condition in which each plant was inoculated with the same bacterial strain. Controls were inoculated with sterile osmosis water. The dishes were rotated every 24 h to avoid boundary effects.

### Barcode and Sanger sequence data analysis

Amplicon sequence variants (ASV) and Sanger sequences were aligned with MUSCLE (Edgar 2004) and manually edited and corrected with Genedoc (73), then the phylogeny was inferred by the neighbour joining method (74) with 1,000 bootstrap analyses for distance calculation on MEGA11 (75). The phylotree was displayed with supplementary information using the online iTOL tool (76).

Cross-comparison of our collection diversity with the metabarcoding dataset of (60) was done using the Ribosomal Database Project (RDP) online ‘Library Compare’ tool (77). Note that Barro’s rice sampling for microbiome characterization was performed during the rice flowering stage.

We assumed that the ASV with 99.5% (no more than two mismatches over 400 bp) to 100% ID with the rRNA 16S sequence could represent the analysed isolate. For 10 rRNA 16S sequences, only one corresponding an ASV sequence with 100% ID was found. In the case of three other isolates, no representative ASV with 100% ID was found, so the ASV with 99% ID was selected (harbouring one or two mismatches of over 400 to 425 bp). The representative ASVs were then analysed for their presence and abundance at the sampled sites.

### Statistical analyses and figures preparation

With regard to the data analysis of *in vitro* and *in planta* PGP capacities tests findings, Excel 2016 was used to record the data and R software was used for statistical analyses. The *in planta* PGP capacity test data were analysed separately for each tray to avoid tray effects. A Wilcoxon non-parametric test was used to compare means of two independent or paired samples and detect significant differences (p=0.05). Pearson correlation tests were performed to determine if the *in vitro* PGP capacities of isolates influenced their interactions with plants (*in planta* tests).

A map of the region with sampling locations displayed was created in QGIS (QGIS.org, %Y. QGIS Geographic Information System. QGIS Association. http://www.qgis.org). For the analysis of the diversity of bacterial genera and functional characterization, bar graphs were constructed in Excel spreadsheets, and rarefaction curves of isolate diversity concerning sampling sites and culture media were generated using the “rarefy” function, available in the R vegan package (78) with script already published (https://github.com/lmoulin34/Article_Moussa_culturingbias/blob/main/Script_Dada2_Phyloseq_Fig2). Venn diagrams showing the number of common and specific genera according to the culture medium were generated online (https://bioinformatics.psb.ugent.be/webtools/Venn/). Figures including the phylogeny trees and hit maps were created with the online iTOL tool (76).

## RESULTS AND DISCUSSION

### Sampling strategy of the rice root-associated collection

As shown in Table 1 and Fig 1, to establish the rice root-associated bacteria collection, root samples were collected in rice fields over three consecutive years (2017 to 2019) at 22 sites in two studied rice growing regions in western Burkina Faso (56,60), and at different rice plant development stages. Phenotypically distinct bacterial colonies obtained from whole roots (total microflora) or from surface sterilised roots (putative endophytes) and subsequently plated on five different culture media (TSA of 10 and 50%, NGN, FNb, Baz, PCAT) were reisolated for further taxonomic and phenotypic characterization. The adopted sampling and isolation strategy aimed to maximise the bacterial isolate diversity, not to analyse the geographical or diachronic bacterial diversity.

As shown in Table 2, 3,855 isolates were obtained from four sites. The highest number of isolates (1,668) came from Bama, i.e. the site sampled during the first two years of the study, with the lowest number from Karfiguela (323). However, total numbers of isolates from the RL and IR systems were comparable (1,864 and 1,991, respectively). Regarding the culture media, the highest number of isolates was obtained with TSA (both TSA10 and TSA50), a generalist medium, and it decreased systematically in the following order: TSA>NGN>NFb>PCAT>Baz. A higher number of isolates was obtained from whole roots (total microflora) than from surface sterilised roots (endophytes). Moreover, a small number of colonies grew on selective PCAT and Baz media, resulting in a limited number of bacteria that could be isolated from these media (Table 2).

**Table 2.** Summary of all sampled isolates and the part constituting the working collection.

| Ecology | Sampl. site | Total bacterial isolates (characterized isolates) |  |  |  |  |  |  |  |  |  | Total |
| --- | --- | --- | --- | --- | --- | --- | --- | --- | --- | --- | --- | --- |
|  |  | TSA tot | TSA end | NGN tot | NGN end | NFb tot | NFb end | PCAT tot | PCAT end | Baz tot | Baz end |  |
| RL | Badala | 264<br>(37) | 137<br>(28) | 207<br>(47) | 76<br>(18) | 183<br>(54) | 58<br>(36) | 46<br>(19) | 21<br>(20) | 0<br>(0) | 0<br>(0) | 992<br>(259) |
| RL | Tengrela | 259<br>(34) | 165<br>(16) | 102<br>(27) | 58<br>(21) | 135<br>(47) | 89<br>(17) | 44<br>(11) | 20<br>(5) | 0<br>(0) | 0<br>(0) | 872<br>(178) |
| IR | Bama | 450<br>(64) | 334<br>(69) | 183<br>(42) | 118<br>(33) | 271<br>(103) | 157<br>(55) | 111<br>(56) | 3<br>(0) | 30<br>(25) | 11<br>(11) | 1668<br>(458) |
| IR | Karfiguela | 90<br>(35) | 104<br>(23) | 23<br>(9) | 15<br>(5) | 66<br>(42) | 24<br>(5) | 1<br>(0) | 0<br>(0) | 0<br>(0) | 0<br>(0) | 323<br>(118) |
| total |  | 1063<br>(169) | 740<br>(136) | 515<br>(125) | 267<br>(77) | 655<br>(246) | 328<br>(113) | 202<br>(86) | 44<br>(25) | 30<br>(25) | 11<br>(11) | 3855<br>(1013) |

Sampl. site, sampling site; root compartment: tot, total root; end, endophytic root.

### Bacterial working collection diversity

Isolate selection was aimed, as best possible, at characterizing a balanced number of isolates on the basis of the sampling location, rice ecology and culture medium used for isolation (Fig 2). The highest number of characterized isolates originated from Bama (458 isolates) where high microbiome diversity was reported by (60) (Supp. Fig. 5), in comparison to those from Badala, Karfiguela and Tengrela (259, 118 and 178 isolates, respectively)(Table 2), which enabled us to constitute a working collection of 1,013 isolates. The characterized isolates are described (sampling year and site, PGP characteristics of the obtained isolates, 16S rRNA sequences and their accession numbers) in S1 Table. As shown by the rarefaction curves at the genus level (Figs 2A and B, S1 Fig), the maximum diversity of the sampled bacteria had not yet been reached in terms of sites and culture media. A comparison with data representing the potential diversity of rice root-associated culturable bacteria from Bama reported by (49) (Supp. Fig S2) suggested that about 60, 45 and 38% of the diversity had been reached for TSA, NFb and NGN, respectively.

**Fig 2.**
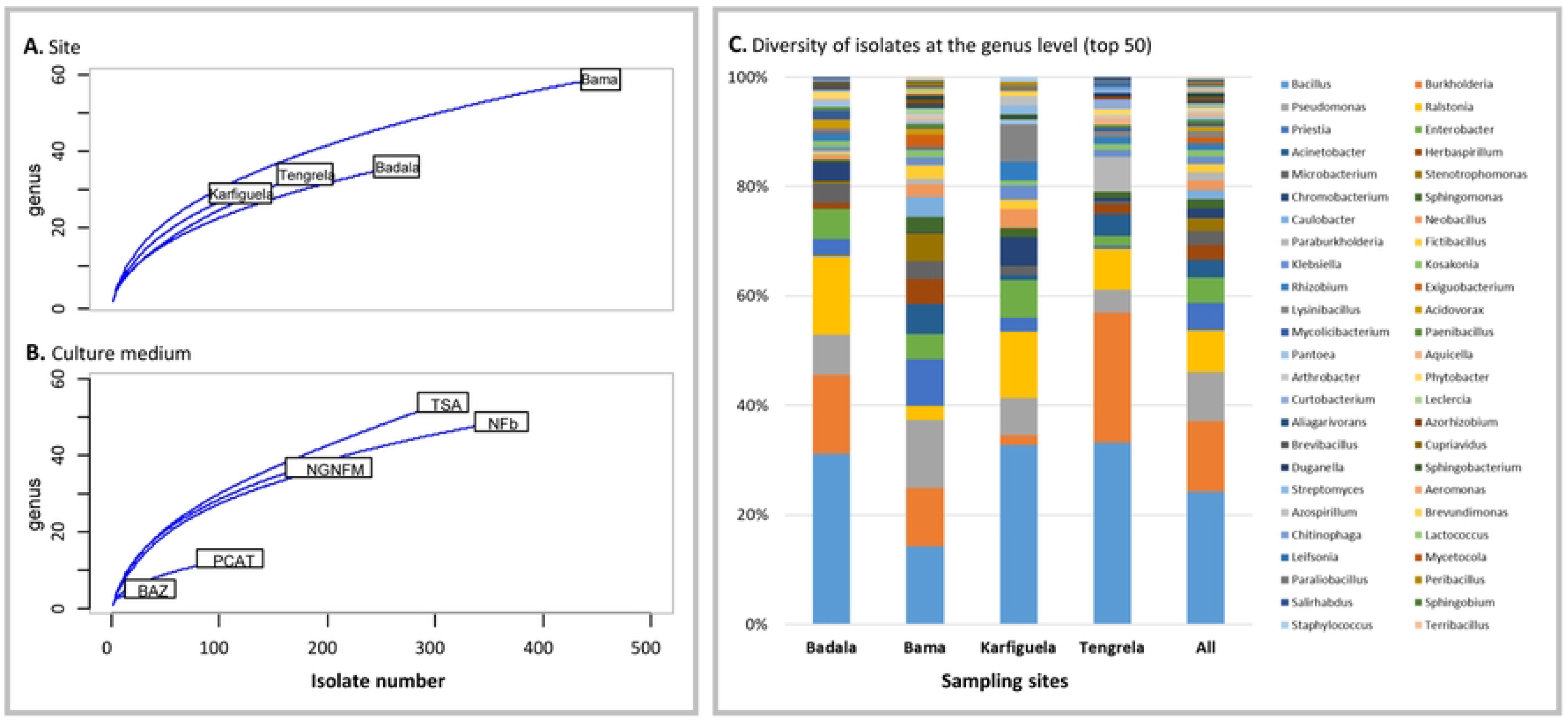
Rarefaction curves (A) and (B), and diversity of isolates obtained from different sites at the genus level (C).

As shown in Fig 2C and S1 Fig the diversity of our working collection covered four bacterial phyla (Actinobacteria, Bacteroidetes, Firmicutes and Proteobacteria, with alpha, beta and gamma subclasses) and encompassed 83 bacterial genera, including *Bacillus* (236 isolates), *Burkholderia* (125), *Pseudomonas* (87), *Ralstonia* (75), *Priestia* (45), *Enterobacter* (46), *Acinetobacter* (36), *Herbaspirillum* (27), *Microbacterium* (26), *Stenotrophomonas* (22), *Chromobacterium* (17), *Sphingomonas* (17), *Caulobacter* (17) and *Paraburkholderia* (16). To the best of our knowledge, two other large collections of rice-associated bacteria have been reported. (38) characterized 689 isolates from a single culture medium (TSA), covering the same four phyla with a higher proportion of Actinobacteria and covering 51 genera (S1 Fig). Higher diversity was achieved by Zhang *et al.* (39) using a high-throughput isolation strategy, and again with only one culture medium (TSA). The final working collection consisted of 1,041 isolates, covering five bacterial phyla (with the fifth phylum being Deinococcus) and 138 genera. Finally, data obtained by Sondo *et al.* (49) via direct barcoding of culturable bacteria grown on five different culture media (cultured bacteria were scraped from the agar plates and used directly for DNA extraction, without further cultivation steps) indicated the presence of bacteria belonging to five phyla (with the fifth phylum being Verrucomicrobia) and to 148 genera. These results suggest that it should be possible to obtain isolates belonging to two more phyla, i.e. Verrucomicrobia and Deinococcus, without changing the culture conditions, but by sequencing all isolates since the corresponding bacteria were found to have a low abundance of 0.4 to 0.6% (S1 Fig).

### Culture medium efficiency for growth of diverse taxa

The taxonomic assignation of isolates obtained with each culture medium is shown in Fig 3A. The isolate diversity comparison revealed that the generalist TSA medium, as well as two selective N_2_-free media (NFb and NGN), allowed the recovery of high isolate diversity covering 54, 45 and 34 genera, respectively (Fig 3A). As shown in the Venn diagram (Fig 3B), TSA shared 26 genera with NGN and 27 with NFb, and 17 genera overall. The same trend, where TSA, NFb and NGN were found to function as efficient culture media, was observed by Sondo et al. (49). Thus, TSA, as well as NFb and NGN, seemed to be the efficient culture media for the diversity study, with each of them increasing the overall diversity by providing their specific part, representing from 7.9% to 22% of the overall diversity (Fig 3B). TSA and NFb maintained the diversity richness more than NGN, allowing the isolation of 16 and 17 specific genera, respectively (including *Acidovorax*, *Gordonia*, *Xanthomonas* for TSA and *Achromobacter*, *Azospirillum*, *Nitrospirillum*, *Sphingobacterium* and *Vogesella* for NFb). Finally, PCAT and BAz media, which were not used by Sondo *et al.* (49), resulted in a low number of colonies and consequently a low isolate diversity (13 and 5 genera, respectively, Fig 3A). PCAT shared 10 bacterial genera with TSA and NFb and 9 with NGN, but only allowed the isolation of one specific genus (*Microbulbifer*). Consequently, PCAT, which has been described as a specific medium for *Burkholderia* isolation (62), would indeed be an interesting medium to isolate a specific genus, but would not help to increase the diversity if TSA, NGN and NFb are already used. BAz had no specific genera as it shared its four genera with all other media and one with TSA.

**Fig 3.**
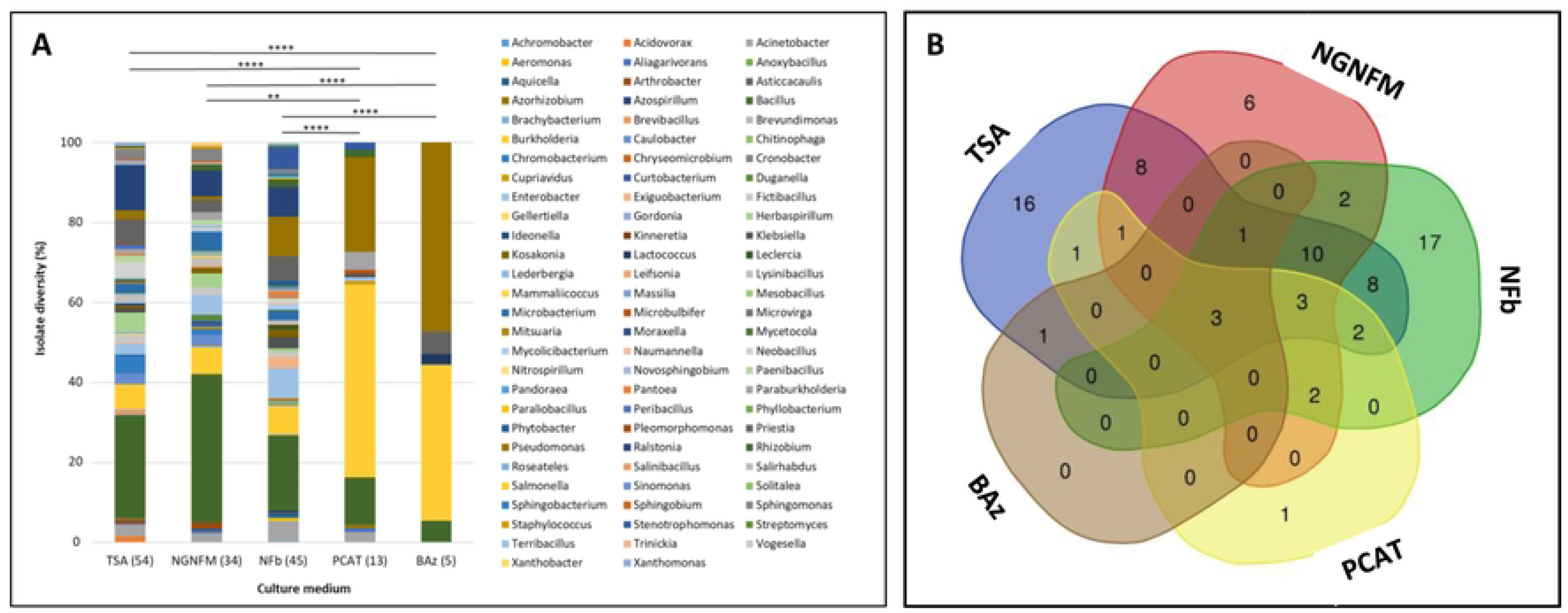
Bacterial isolate diversity sampled from each culture medium. A, barplot representing the number of genera (in brackets) per culture medium. B, Venn diagram showing the number of common and specific genera according to the culture medium.

### Characterization of functional *in vitro* PGP capacities

All isolates from the working collection were screened *in vitro* for four plant growth promotion capacities: ammonia production (AP), auxin biosynthesis capacity (ABC), siderophore biosynthesis capacity (SBC) and phosphate solubilising capacity (PSC). PGP capacities were generally analysed as features of individual isolates in order to be able to predict their potential PGP capacities, and/or to study direct plant-bacterium interactions (30,79). However, we have to keep in mind that the plant root system is immerged in (and inhabited by) the microbial population, whose members harbour specific and shared PGP capacities, and thus the plant interacts with a population of taxonomically different bacteria and a pool of synthetized molecules. Thus, the analysis of four PGP capacities of over 1,000 isolates allows the global view on the root associated bacterial population presented in the hitmap (Fig 4A) and in the principal component analysis bi-plot (Fig 4B), with the general trends showcased on the basis of the bacterial taxonomy. Almost all isolates from four bacterial phyla showed an ability to produce ammonia, with only 6.9% of isolates being negative. ABC was the second the most abundant PGP capacity and was detected in at least half of the Actinobacteria, Bacteroidetes, alpha and gamma Proteobacteria isolates. As suggested by the principal component analysis findings, ABC appeared to be negatively correlated to PSC, the less frequent among isolates (Figs 4A and B). In this regard, Actinobacteria and Gammaproteobacteria seemed to be enriched in ABC (57 and 67% of isolates, respectively), Betaproteobacteria in SBC (51%) and PSC (39%), while Firmicutes displayed moderate-low ABC, PSC and SBC capacities (27, 25.6 and 9%, respectively). As shown in Fig 4C, some genera and species in four phyla seemed to have a high number of PGP capacities. There was a noteworthy presence of two species belonging to two phyla having the lowest number of isolates in the working collection, i.e. *Microbacterium azadirachtae* in the Actinobacteria phylum and *Sphingobacterium multivorum* in the Bacteroidetes phylum. In the Firmicutes phylum, species belonging to the *Bacillus* and *Priestia* genera (*Bacillus aerius*, *B. pumilus*, *B. tropicus* and *P. aryabhattai*) seemed to stand out in terms of the accumulation of PGP capacities. Finally, in the Proteobacteria phylum, remarkable species were noted in the following genera: *Rhizobium* (*R. leucaenae*, *R. straminoryzae*), *Burkholderia* (*B.ambifaria*, *B. anthina*, *B. cepacia*, *B. contaminans*, *B. latens*, *B. pyrrocinia*, *B. stagnalis*, *B. vietnamiensis*), *Paraburkholderia* (*P. kururiensis*), *Ralstonia* (*R. mannitolilytica*, *R. picketti*), *Aquitalea* (*A. pelogenes*), *Herbaspirillum* (*H. aquaticum*, *H. seropedicae*), *Chromobacterium* (*C. violaceum*), *Vogesella* (*V. urethralis*), *Acinetobacter* (*A. lactucae*, *A. seifertii, A. nosocomialis*) and *Pseudomonas* (*P. guezennei*, *P. monteilii*, *P. panipatensis*, *P. taiwanensis*). Note, however, that despite the overall trends, high variability between isolates was observed even within a single species (Fig 4D). This observation, brings the question of the significance of a single isolate or even a single species and it’s PGP capacities in the context of the bacterial population. For example, indole-3-acetic acid (IAA), i.e. the most crucial and common natural auxin which, following production by bacteria, stimulates plant root hair formation, increases the number and length of lateral and primary roots (when it occurs within an ideal concentration range), but also inhibits primary root growth at higher concentrations (reviewed by (80)). Our findings revealed that the IAA biosynthesis capacity seemed to be widespread among culturable rhizobacteria. Similar results were obtained by Zhang *et al*. (17) who analysed 7,282 prokaryotic genomes and showed that 82.2% of the corresponding bacteria should potentially be capable of synthesizing IAA from tryptophan or intermediates. There is still little knowledge about how a plant interacts with a bacterial population that harbours abundant IAA biosynthesis capacities, and also how diverse bacterial taxa regulate the expression of these capacities. The availability of a pool of isolates belonging to the same species, and/or also those from different species and taxonomic groups, will facilitate further studies.

**Fig 4.**
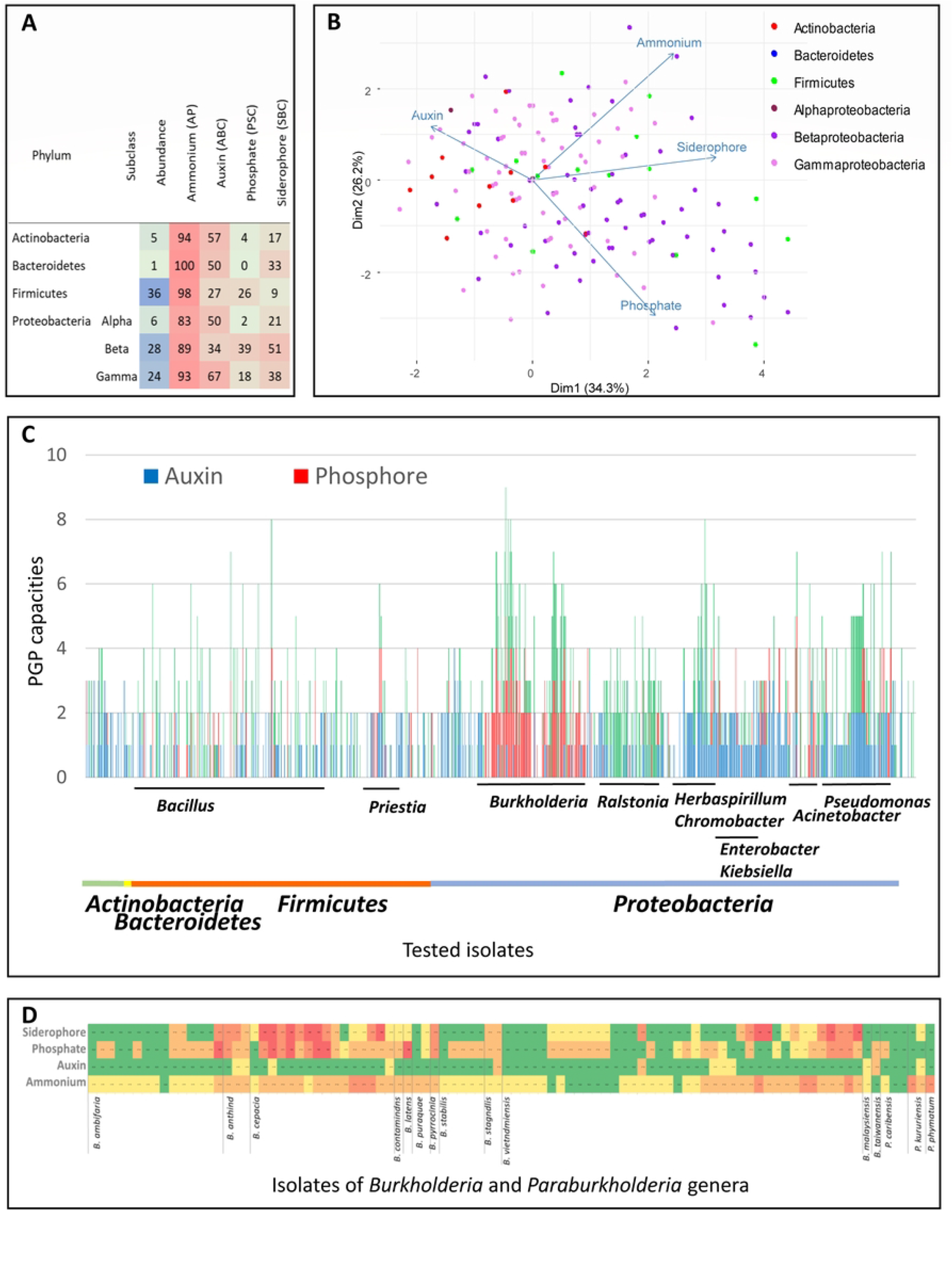
Characterization of PGP capacities of isolates in the working collection. A, proportion of isolates (in percentage) presenting PGP capacities on the basis of the bacterial phyla: Ammonium, ammonia production; Auxin, auxin biosynthesis capacity; Phosphate, phosphate solubilising capacity and Siderophore, siderophore biosynthesis capacity. B, principal component analysis. C, detailed representation of *in vitro* PGP capacities (y-axis) of the working collection isolates (x-axis) on the basis of the taxonomy. As a majority of isolates had an ammonia production capacity, it was not displayed in part C of the figure.

### Bacterial inoculation effect on early plant growth

We screened our bacterial collection for plant growth promotion effects on early plant developmental stage by selecting 45 isolates (S1 and S2 Tables), representative of the diversity of the collection, from Actinobacteria, Bacteroidetes, Firmicutes and Proteobacteria phyla for *in planta* tests. The plant-bacteria interaction was assessed on two rice cultivars (TS2 and Orylux 6) commonly grown and consumed in Burkina Faso (which also served as plant hosts for some isolates), both belonging to two different and well separated clades within *Oryza sativa indica*, as reported by Barro et al. (53). A broad range of plant responses was observed two weeks after bacterial inoculation, ranging from a growth-promoting effect to a negative effect on plant development (Fig 5, S2 Table). Furthermore, the effects of a given bacterium varied according to the plant genotype. Only five isolates showed a significant PGP effect on both rice cultivars. These isolates belonged to the Proteobacteria phylum, i.e. *Kosakonia oryzendophytica* ABIP 2271, *Aquicella siphonis* ABIP 3002, and *Caulobacter vibrioides* ABIP 1460, and Firmicutes phylum, i.e. *Bacillus pumilus* ABIP 2917 and *Fictibacillus rigui* ABIP 1663. For TS2, the maximum increase in leaf weight (LW) and root weight (RW) was observed with *Caulobacter vibrioides* ABIP 1460 (43 and 37%) and, in the case of Orylux 6, *Bacillus pumilus* ABIP 2917 gave the best results (79 and 101%) (Fig 5).

**Fig 5.**
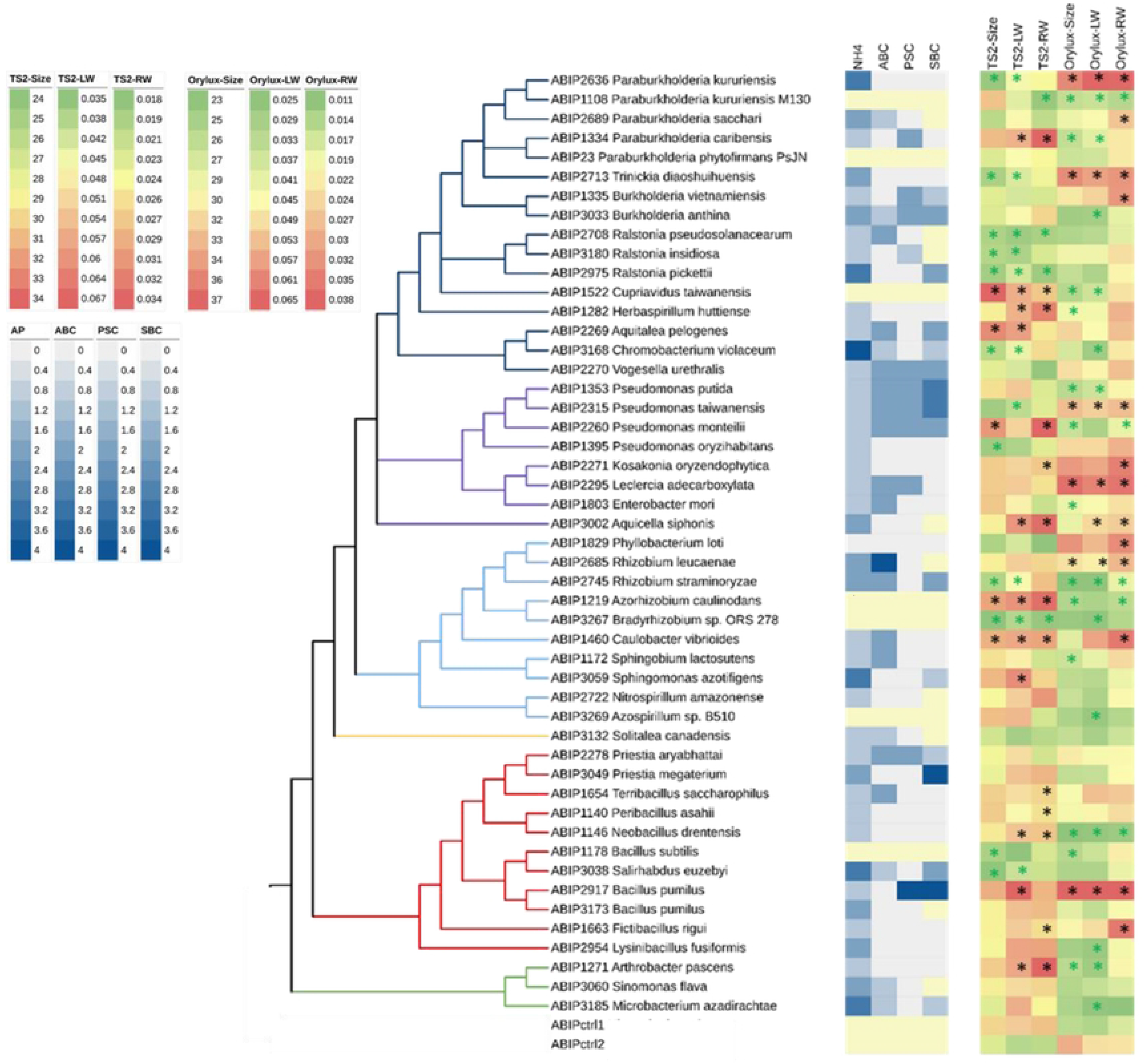
Comparison of *in vitro* measured PGP capacities and *in planta* PGP effects of 45 representative isolates according to their taxonomy. Asterisks indicate statistically significant PGP inoculation effects-black for positive and green for negative effects. NH4, ammonium; ABC, auxin biosynthesis capacity; PSC, phosphate solubilisation capacity; SBC, siderophore biosynthesis capacity; TS2 and ORYLUX, rice cultivars, ctrl1 and ctrl2, for controls with non-inoculated plants; Size, plant size (cm); LW, aerial part dry weight (g); RW, root dry weight (g).

Five isolates had a PGP effect on one cultivar and no significant effect on the second. For TS2, this effect was observed with *Aquitalea pelogenes* ABIP 2269, *Terribacillus saccharophilus* ABIP 1654 and *Peribacillus asahii* ABIP 114, while for Orylux 6 the effect was noted with *Rhizobium leucaenae* ABIP 268 and *Leclercia adecarboxylat* ABIP 2295. However, several isolates that had a positive effect on one genotype had significantly opposite effects on the second rice genotype. For instance, for seven isolates (*Paraburkholderia caribensis* ABIP 1334, *Cupriavidus taiwanensis* ABIP 1522, *Herbaspirillum huttiense* ABIP 1282, *Pseudomonas monteilii* ABIP 2260, *Azorhizobium caulinodans* ABIP 1219, *Neobacillus drentensis* ABIP 1146, *Arthrobacter pascens* ABIP 1271), a significant PGP effect was observed on TS2, yet a decrease in leaf size or root weight was observed on Orylux 6. Conversely, three isolates (*Paraburkholderia kururiensis* ABIP 2636, *Trinickia diaoshuihuensis* ABIP 2713, *Pseudomonas taiwanensis* ABIP 2315) showed a PGP effect on Orylux 6 but reduced TS2 development. Among isolates showing the positive effects on seedlings growth, the species, *Bacillus pumilus Caulobacter vibrioides* and *Kosakonia oryzendophytica* (previously *Enterobacter oryzendophytica* (81)) have been already reported as able to improve the rice growth (38,79,82). In contrast, as far as we know, this is the first study that reports on interaction with rice and their capacities to improve seedlings growth for seven species *Aquicella siphonis*, *Aquitalea pelogenes, Fictibacillus rigui*, *Leclercia adecarboxylata* (83), *Peribacillus asahii* (*Bacillus asahii)* (85)(86), *Rhizobium leucaenae* (87) and *Terribacillus saccharophilus* (84). Finally, *Chromobacterium violaceum* ABIP 3168, *Rhizobium straminoryzae* ABIP 2745 and *Bacillus subtilis* ABIP 1178 (and also *Paraburkholderia kururiensis* ABIP 1108 and *Bradyrhizobium* sp. (ORS 278) ABIP 3267, i.e. the reference isolates used in the study) showed significant negative effects on both rice genotypes. Overall, the most significant effects of the LW increase were obtained in the case of TS2 with *Azorhizobium caulinodans* ABIP 1219 (47.2%), *Caulobacter vibrioides* ABIP 1460 (43%) and *Arthrobacter pascens* ABIP 1271 (42%), and in the case of Orylux 6 with *Pseudomonas taiwanensis* ABIP 2315 (157.7%), *Paraburkholderia kururiensis* ABIP 2636 (104.2%) and *Leclercia adecarboxylata* ABIP 2295 (88.2%) (S2 Table).

Statistical analysis of the *in vitro* and *in planta* results revealed that the PGP bacterial capacity and PGP effect on seedlings were not correlated, regardless of the rice genotype (S2 Table). Furthermore, we observed a specific interaction between bacteria and the rice genotype. Finally, high variability in the plant-bacteria interaction was observed even within the same bacterial species, as illustrated by the following example of isolates belonging to *P. kururiensis* and *B. pumilus* species. Isolates of *P. kururiensis*, ABIP 1108 (isolate M130 from Brazil), which has been reported to have strong growth-promoting effects and an ability to increase rice yields (88,89), and ABIP 2636 (from Burkina Faso, this study) had different effects on the tested rice cultivars, suggesting bacterial adaptation to a specific rice genotype and probably to specific climatic and environmental conditions, as discussed by Liu, Qin and Bai (22). In the case of *B. pumilus*, ABIP 2917 had a marked significant PGP effect on both rice cultivars, while ABIP 3173 showed no effect. Genotypic, and thus phenotypic, variability within the same species is common among bacteria and has been observed even for isolates from different compartments within the same plant (90).

### Comparison of isolates versus amplicon barcode diversity

We assessed the extent of representativeness of our bacterial collection in terms of a potential SynCom design by comparing a pool of rRNA 16S sequences with 16S amplicon barcode ASV data (cut-off at 0.0001 abundance) obtained in the study of Barro *et al.* (60), on the same rice fields, rhizosphere soil and roots (60). At the phylum level, our collection covered four (19%) of 21 phyla, 19 (32.8%) of 58 orders and finally 70 (33%) of 212 genera detected in the barcode data (Fig 6, S2 Table). Note that the barcode diversity data derived from rice plants sampled during flowering, while the sampling in this study was done during different rice developmental stages. Although the total diversity represented between 19 and 33% of the barcode diversity at different taxonomic levels, it covered the most abundant taxonomic clades within the barcode diversity of Proteobacteria, Firmicutes and Bacteroidetes phyla (Fig 6). It also included potential representatives of the core microbiome and microbiome hub species identified in the barcode data (60), which belonged to seven of the nine identified genera (core microbiome: *Paraburkholderia*, *Ralstonia*; and hubs: *Bacillus*, *Neobacillus*, *Enterobacter*, *Aeromonas*, *Acinetobacter*). Previous studies on rice-associated bacteria (38,39) and barcoding of culturable bacteria (49) suggest that it should also be possible to isolate the taxa from Deinococcus and Verrucomicrobia phyla without changing the culture conditions and to improve the diversity within the four remaining phyla, especially Bacteroidetes (three genera represented by ABIP isolates compared to 14 genera of different ASVs) and Proteobacteria (46 genera of isolates compared to 76 genera of ASVs, including Deltaproteobacteria) as shown by Sondo *et al.* (49). However, there is still a major share of missing microbiome diversity potentially influencing plant development and health, although its abundance appears to be lower and scarce for some genera. Indeed, several studies have highlighted the role of rare species (also called satellite taxa) in plant-microbe interactions and, more generally, in key ecosystem functions (91,92). Therefore, in addition to the application of new conditions with plant-based culture media, as reviewed by Sarhan *et al.* (45), accurate knowledge on missing taxa could facilitate our search for specific culture conditions. First, like other bacterial collections, ours lacked the anaerobic bacteria that thrive in the rice roots and rhizosphere, as observed by Edwards et al. (6). These bacteria have been extensively studied for their contribution to anaerobic degradation of plant polymers (40) and iron reduction (93,94), but they have never been included in the rice/microbiome interaction experiments. Nevertheless, numerous anaerobic taxa observed in our barcoding data, such as those belonging to the Clostridiaceae1 family in the Firmicutes phylum, the Opitutaceae family and Verrucomicrobia subdivision3 in the Verrucomicrobia phylum, the Geobacteraceae family or the Desulfovibrionaceae family in the Deltaproteobacteria class, could be obtained as culturable isolates from rice fields, as previously reported (95–98), so it should be possible to include them in the SynCom experiment. Second, we also searched for taxa missing from our collection, while taking the diversity (number of ASVs) and abundance (number of reads per ASV) of ASVs identified per genus into account. The major missing phylogroups within Proteobacteria, which were generally well represented in our collection, thus included the entire Deltaproteobacteria class and, within the Gammaproteobacteria class, two of Betaproteobacteriales families (Nitrosomonadaceae, Rhodocyclaceae). Within the Actinobacteria phylum, which was not well represented in our collection (Fig 7), several orders were missing, with the most important being Micromonosporales, Acidimicrobiales and Kineosporiales, and within the Bacteroidetes phylum the missing orders were Bacteroidales, Flavobacteriales and Cytophagales. Finally, Acidobacteria, Chloroflexi phyla, which were abundant in the barcode diversity, Patescibacteria, Spirochaetes, Armatimonadetes, Verrucomicrobia and Nitrospirae, which were less abundant, and finally the scarce phyla Fusobacteria, Gemmatimonadetes, Latescibacteria, Rokubacteria, WSP-2 and Epsilonbacteraeota were completely missing (Fig 7). With the exception of the Patescibacteria, Fusobacteria, Latescibacteria, Rokubacteria and WSP-2 phyla, which are currently candidate phyla lacking cultured representatives (99–103), viable isolates could be obtained for other mentioned phyla. For example, the culture conditions for isolating bacteria from Acidobacteria, Verrucomicrobia (104–106) and Spirochaetes phyla (107,108) have been described. New strategies based on the prediction of new organism-medium combinations using already existing media (KOMODO online tool, (109)) or using metagenome-available data should also help in modelling specific media for specific taxa, as described by Liu *et al.* (110).

**Fig 6.**
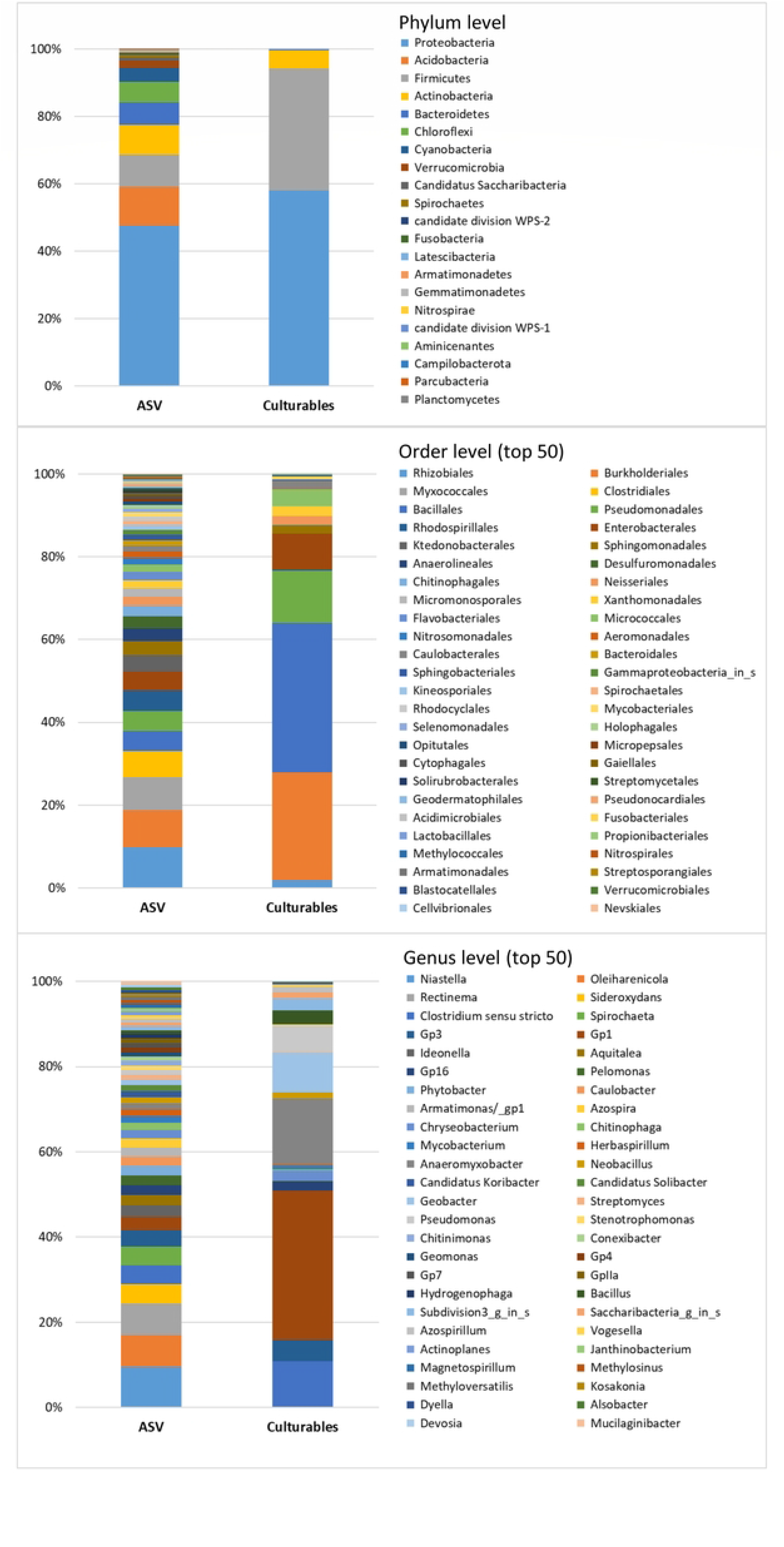
Comparison of the diversity of isolates between the culturable collection and 16S amplicon barcode data from the study of Barro *et al.* (2022) with a cut-off at 0.0001%. A, phylum level; B, top 50 orders; C, top 50 genera.

**Fig 7.**
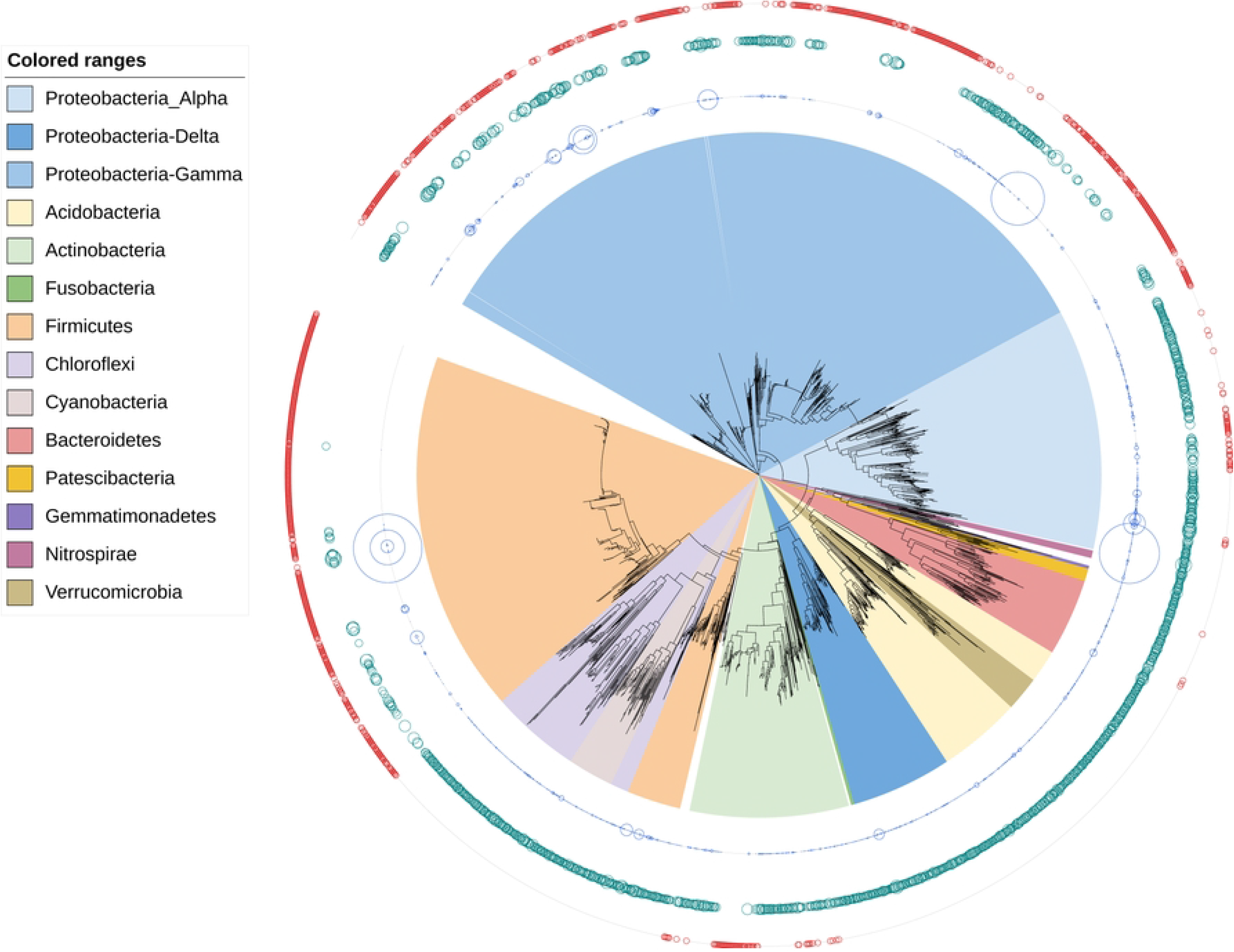
Neighbour joining phylogenetic tree of combined rice bacterial associated microbiota diversity (ASVs and culturable isolates) The collection isolates are indicated by red circles. ASVs abundance is displayed with circles (blue circles, 1/100 read number (to better highlight the most abundant ones); green circles, log of read number).

### PGPR occurrence in the rice root microbiote

Representative ASVs were found for 13 isolates and their presence in the rice root/rhizosphere microbiome is shown in Fig 8. Among the ASVs representing isolates with PGP capacities, five appeared to be very abundant, with a total of 4,000 to 48,400 reads each, three were less abundant, with a total of 1,000 to 2,000 reads each, while the remaining five ASVs were scarce, with a total of 160 to 700 reads (Fig 8). Finally, a representative ASV for *A. siphonis* ABIP 3002 could not be identified in the barcode data (although we searched the entire dataset, including before the cut-off). The fact that we could not detect an ASV with > 95% ID within an overlapping 425 bp sequence, suggests its very low abundance in the root/rhizosphere microbiome.

**Fig 8.**
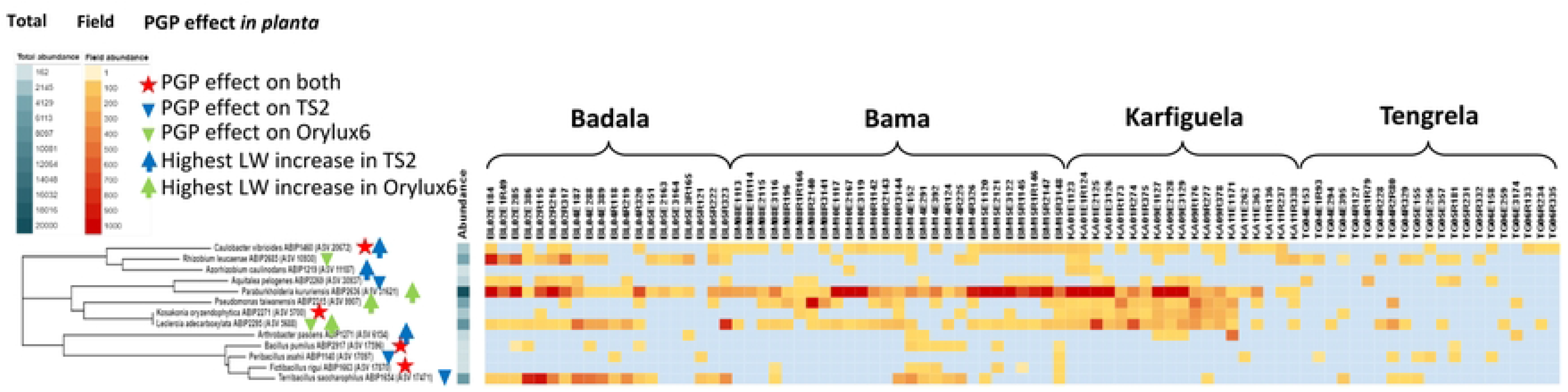
Assessment of PGPR in the rice root microbiome across the studied fields based on a comparison with the corresponding ASVs. Irrigated conditions, Bama and Karfiguela; and rainfed lowland conditions, Badala and Tengrela. The colour scale represents the number of reads of each ASV.

Interestingly, five isolates with PGP effects on both cultivars were not among the most abundant ASVs analysed. They turned out to be among the rare or less abundant ASVs. However, by far the most abundant ASV appeared to be ASV 31621, representing *Paraburkholderia kururiensis* (ABIP 2636), which only affected Orylux 6, and the three other most abundant ASVs represented isolates also affecting Orylux 6. Moreover, the ASV representing *Paraburkholderia kururiensis* (ABIP 2636) was detected in most of the fields from Bama, Badala and Karfiguela, but was completely absent from the Tengrela site. Two rare ASVs, representing *F. rigui* ABIP 1663 and *A. caulinodans* ABIP 1219, were also absent from Tengrela, and only three ASVs were detected in at least 6 out of 21 fields. These were two abundant isolates, i.e. *Leclercia adecarboxylata* ABIP 2295 and *R. leucaenae* ABIP 2685, and the rare *P. asahii* ABIP 1140, which was almost absent from three other sites. Fields of the Tengrela site showed considerable heterogeneity in the presence/absence of the 13 ASVs analysed. Their presence was not as conserved as in Bama, Badala and Karfiguela. In fact, Tengrela has been described in Barro *et al.* (60) as being a site where rice plants host microbiomes that are statistically different from those of other sites with lower bacterial diversity, possibly due to the presence of different rice cultivars (53) and different agricultural practices (56). ASVs representing the remaining three abundant isolates *R. leucaenae* ABIP 2685 and *T. saccharophilus* ABIP 1654 and *L. adecarboxylata* ABIP 2295 were the most abundant in Badala.

Nevertheless, among the PGPR studied, eight appeared to be abundant in the plant microbiome (number of reads per field) and widely distributed in the rice growing areas, including Tengrela, specific site for its cultural practices and rice cultivars (60). To the best of our knowledge, this is the first study to report the actual presence of screened PGPR in the field.

## CONCLUSION

Rice is a staple food for half the world’s population. In Africa, and particularly in Burkina Faso, its consumption has been increasing since 1980s as a result of demographic trends, rural-urban migration and changing lifestyles. Despite the increase in the area under rice cultivation (FAOSTAT, 2023; https://fenix.fao.org/faostat/internal/fr/#data/QC), Burkina Faso still has to import 57% of the rice consumed in the country (111). Plant root-associated microorganisms—due to their application in crop production and protection—have attracted increasing attention (112). In Burkina Faso, studies have been initiated on the diversity of rice rhizosphere and root microbiomes using metabarcoding (60) and culturable bacteria (49) approaches to analyse the effects of rice production systems (irrigated cropping areas vs rainfed lowland conditions) on microbiome diversity.

With the aim of developing future agroecological practices based on microorganisms available *in situ* (92), which seem to be of particular importance in developing countries, through their direct use as inoculants or via the adaptation of ‘microbiome-friendly’ cultural practices, efforts have been made to systematically isolate bacteria associated with rice roots. The present paper describes a bacterial collection whose taxonomic diversity covers 33% of barcoded microbiome diversity at the genus level, including core microbiome and potential microbiome hub representative bacteria, essential for future SynCom functional studies. Cross-referencing with barcode data indicates which new taxonomic groups we should be looking for, which should thus help to adjust culture conditions to progress in the process of the missing taxa isolation. Our results revealed that, among the isolates with PGP capacity (on seedlings), several appeared to be significant in terms of bacterial abundance in the root microbiome, while others were observed at much lower levels, so these could be excellent candidates for functional studies. With a view to using PGPR as inoculants in the field, these results highlight the need to carry out comparative tests on the effectiveness of using abundant vs. scarce PGPR to understand the reasons for (e.g. soil characteristics) and consequences of (effects on plants) differences in abundance and also the efficiency of their survival as inoculated bacteria.

## Acknowledgments

The authors thank Abdoul Kader Guigma, Sylvain Zoungrana, Issouf Sanga, Manaka Douanio, Daouda Hema, Abalo Raoul Kassankogno, Gilles Béna, Didier Tharreau and Christophe Brugidou for their contributions to the field sampling in Burkina Faso and transport of samples to France. We thank the rice farmers at the Badala, Bama, Tengrela, and Karfiguela sites for their kind collaboration.

## Funding

This work was publicly funded by the CGIAR Research Program on Rice Agri-food Systems (RICE). We thank the French National Research Institute for Sustainable Development (IRD) for awarding MS a PhD Research grant through its ARTS program.

## Author contributions

A.K., L.M. conceived the idea and designed the experiment. A.K., M.S., I.R. conducted the experiments, C.T. and M.B. conducted the field work, A.K., W.I., L.M. and K.K. supervised the global investigation and methodology, W.I. and K.K. were involved in the project administration, A.K. and M.S. analysed the data and wrote the manuscript with input from all other authors.

## Declaration of competing interest

The authors declare that they have no competing interests.

## Notes

### Competing Interest Statement

The authors have declared no competing interest.

## References

1. Bulgarelli D, Schlaeppi K, Spaepen S, van Themaat EVL, Schulze-Lefert P. Structure and Functions of the Bacterial Microbiota of Plants. Annu Rev Plant Biol. 2013 Apr 29;64(1):807–38.

2. Hardoim PR, van Overbeek LS, Berg G, Pirttilä AM, Compant S, Campisano A, et al. The Hidden World within Plants: Ecological and Evolutionary Considerations for Defining Functioning of Microbial Endophytes. Microbiol Mol Biol Rev. 2015 Sep;79(3):293–320.

3. Bai B, Liu W, Qiu X, Zhang J, Zhang J, Bai Y. The root microbiome: Community assembly and its contributions to plant fitness. Integrative Plant Biology. 2022 Jan 13;jipb.13226.

4. Bulgarelli D, Rott M, Schlaeppi K, Ver Loren van Themaat E, Ahmadinejad N, Assenza F, et al. Revealing structure and assembly cues for Arabidopsis root-inhabiting bacterial microbiota. Nature. 2012 Aug;488(7409):91–5.

5. Beckers B, Op De Beeck M, Weyens N, Boerjan W, Vangronsveld J. Structural variability and niche differentiation in the rhizosphere and endosphere bacterial microbiome of field-grown poplar trees. Microbiome. 2017 Dec;5(1):25.

6. Edwards J, Johnson C, Santos-Medellín C, Lurie E, Podishetty NK, Bhatnagar S, et al. Structure, variation, and assembly of the root-associated microbiomes of rice. Proc Natl Acad Sci USA [Internet]. 2015 Feb 24 [cited 2022 Jul 29];112(8). Available from: https://pnas.org/doi/full/10.1073/pnas.1414592112

7. Lee SA, Park J, Chu B, Kim JM, Joa JH, Sang MK, et al. Comparative analysis of bacterial diversity in the rhizosphere of tomato by culture-dependent and -independent approaches. J Microbiol. 2016 Dec;54(12):823–31.

8. Hartman K, van der Heijden MG, Roussely-Provent V, Walser JC, Schlaeppi K. Deciphering composition and function of the root microbiome of a legume plant. Microbiome. 2017 Dec;5(1):2.

9. Walters WA, Jin Z, Youngblut N, Wallace JG, Sutter J, Zhang W, et al. Large-scale replicated field study of maize rhizosphere identifies heritable microbes. Proc Natl Acad Sci USA. 2018 Jul 10;115(28):7368–73.

10. Santos-Medellín C, Edwards J, Liechty Z, Nguyen B, Sundaresan V. Drought Stress Results in a Compartment-Specific Restructuring of the Rice Root-Associated Microbiomes. Ausubel FM, editor. mBio. 2017 Sep 6;8(4):e00764–17.

11. Lundberg DS, Lebeis SL, Paredes SH, Yourstone S, Gehring J, Malfatti S, et al. Defining the core Arabidopsis thaliana root microbiome. Nature. 2012 Aug;488(7409):86–90.

12. Agler MT, Ruhe J, Kroll S, Morhenn C, Kim ST, Weigel D, et al. Microbial Hub Taxa Link Host and Abiotic Factors to Plant Microbiome Variation. Waldor MK, editor. PLoS Biol. 2016 Jan 20;14(1):e1002352.

13. Vannier N, Agler M, Hacquard S. Microbiota-mediated disease resistance in plants. Zipfel C, editor. PLoS Pathog. 2019 Jun 13;15(6):e1007740.

14. Douglas GM, Maffei VJ, Zaneveld JR, Yurgel SN, Brown JR, Taylor CM, et al. PICRUSt2 for prediction of metagenome functions. Nat Biotechnol. 2020 Jun;38(6):685–8.

15. Sessitsch A, Hardoim P, Döring J, Weilharter A, Krause A, Woyke T, et al. Functional Characteristics of an Endophyte Community Colonizing Rice Roots as Revealed by Metagenomic Analysis. MPMI. 2012 Jan;25(1):28–36.

16. 16. Howe A, Yang F, Williams RJ, Meyer F, Hofmockel KS. Identification of the Core Set of Carbon-Associated Genes in a Bioenergy Grassland Soil. Kelly JJ, editor. PLoS ONE. 2016 Nov 17;11(11):e0166578.

17. Zhang P, Jin T, Sahu S Kumar, Xu J, Shi Q, Liu H, et al. The Distribution of Tryptophan-Dependent Indole-3-Acetic Acid Synthesis Pathways in Bacteria Unraveled by Large-Scale Genomic Analysis. Molecules. 2019 Apr;24(7):1–14.

18. Doolittle WF, Zhaxybayeva O. Metagenomics and the Units of Biological Organization. BioScience. 2010 Feb;60(2):102–12.

19. Jansson JK, Hofmockel KS. The soil microbiome — from metagenomics to metaphenomics. Current Opinion in Microbiology. 2018 Jun;43:162–8.

20. Xu L, Pierroz G, Wipf HML, Gao C, Taylor JW, Lemaux PG, et al. Holo-omics for deciphering plant-microbiome interactions. Microbiome. 2021 Mar 24;9(1):69.

21. Beattie GA. Curating communities from plants. Nature. 2015 Dec;528(7582):340–1.

22. Liu YX, Qin Y, Bai Y. Reductionist synthetic community approaches in root microbiome research. Current Opinion in Microbiology. 2019 Jun;49:97–102.

23. de Souza RSC, Armanhi JSL, Arruda P. From Microbiome to Traits: Designing Synthetic Microbial Communities for Improved Crop Resiliency. Front Plant Sci. 2020 Aug 27;11:1179.

24. Ding LJ, Cui HL, Nie SA, Long XE, Duan GL, Zhu YG. Microbiomes inhabiting rice roots and rhizosphere. FEMS Microbiology Ecology [Internet]. 2019 May 1 [cited 2022 Jul 24];95(5). Available from: https://academic.oup.com/femsec/article/doi/10.1093/femsec/fiz040/5420819

25. Adhikari TB, Joseph CM, Yang G, Phillips DA, Nelson LM. Evaluation of bacteria isolated from rice for plant growth promotion and biological control of seedling disease of rice. Can J Microbiol. 2001;916–24.

26. Ferrando L, Mañay JF, Scavino AF. Molecular and culture-dependent analyses revealed similarities in the endophytic bacterial community composition of leaves from three rice (Oryza sativa) varieties. FEMS Microbiol Ecol. 2012 Jun;80(3):696–708.

27. Panhwar QA. Isolation and characterization of phosphate-solubilizing bacteria from aerobic rice. Afr J Biotechnol [Internet]. 2012 Feb 7 [cited 2022 Nov 3];11(11). Available from: http://www.academicjournals.org/AJB/abstracts/abs2012/7Feb/Panhwar%20et%20al.ht m

28. Ji SH, Gururani MA, Chun SC. Isolation and characterization of plant growth promoting endophytic diazotrophic bacteria from Korean rice cultivars. Microbiological Research. 2014 Jan;169(1):83–98.

29. Bai Y, Müller DB, Srinivas G, Garrido-Oter R, Potthoff E, Rott M, et al. Functional overlap of the Arabidopsis leaf and root microbiota. Nature. 2015 Dec;528(7582):364–9.

30. Rolli E, Marasco R, Vigani G, Ettoumi B, Mapelli F, Deangelis ML, et al. Improved plant resistance to drought is promoted by the root-associated microbiome as a water stress-dependent trait: Root bacteria protect plants from drought. Environ Microbiol. 2015 Feb;17(2):316–31.

31. Armanhi JSL, de Souza RSC, de Araújo LM, Okura VK, Mieczkowski P, Imperial J, et al. Multiplex amplicon sequencing for microbe identification in community-based culture collections. Sci Rep. 2016 Jul 12;6(1):29543.

32. Venkatachalam S, Ranjan K, Prasanna R, Ramakrishnan B, Thapa S, Kanchan A. Diversity and functional traits of culturable microbiome members, including cyanobacteria in the rice phyllosphere. Papen H, editor. Plant Biol J. 2016 Jul;18(4):627–37.

33. Samad A, Trognitz F, Compant S, Antonielli L, Sessitsch A. Shared and host-specific microbiome diversity and functioning of grapevine and accompanying weed plants: Microbial communities associated with grapevine and vineyard weeds. Environ Microbiol. 2017 Apr;19(4):1407–24.

34. Borah M, Das S, Baruah H, Boro RC, Barooah M. Diversity of Culturable Endophytic bacteria from Wild and Cultivated Rice showed potential Plant Growth Promoting activities [Internet]. Microbiology; 2018 Apr [cited 2023 Jan 5]. Available from: http://biorxiv.org/lookup/doi/10.1101/310797

35. Levy A, Salas Gonzalez I, Mittelviefhaus M, Clingenpeel S, Herrera Paredes S, Miao J, et al. Genomic features of bacterial adaptation toplants. Nat Genet. 2018 Jan;50(1):138– 50.

36. Moronta-Barrios F, Gionechetti F, Pallavicini A, Marys E, Venturi V. Bacterial Microbiota of Rice Roots: 16S-Based Taxonomic Profiling of Endophytic and Rhizospheric Diversity, Endophytes Isolation and Simplified Endophytic Community. Microorganisms. 2018 Feb 11;6(1):14.

37. Musonerimana S, Bez C, Habarugira G, Bigirimana J, Venturi V. Characterization of bacterial strains from bacterial culture collection of rice sheath in Burundi highlights an Alcaligenes species strain with antibacterial activity against Pseudomonas fuscovaginae rice pathogen. Afr J Microbiol Res. 2021 Oct 31;15(10):497–511.

38. Bertani I, Abbruscato P, Piffanelli P, Subramoni S, Venturi V. Rice bacterial endophytes: isolation of a collection, identification of beneficial strains and microbiome analysis: Beneficial bacterial endophytes of rice. Environmental Microbiology Reports. 2016 Jun;8(3):388–98.

39. Zhang J, Liu YX, Zhang N, Hu B, Jin T, Xu H, et al. NRT1.1B is associated with root microbiota composition and nitrogen use in field-grown rice. Nat Biotechnol. 2019 Jun;37(6):676–84.

40. Liesack W, Schnell S, Revsbech NP. Microbiology of flooded rice paddies. FEMS Microbiol Rev. 2000 Dec;24(5):625–45.

41. Phung LD, Miyazawa M, Pham DV, Nishiyama M, Masuda S, Takakai F, et al. Methane mitigation is associated with reduced abundance of methanogenic and methanotrophic communities in paddy soils continuously sub-irrigated with treated wastewater. Sci Rep. 2021 Apr 1;11(1):7426.

42. Otoidobiga CH, Kam H, Bagayogo A, Savadogo A, Sawadogo JB, Sawadogo S, et al. Effect of Combined Application of Subsurface Drainage and Mineral Fertilization on Iron-Reducing Bacterial Populations’ Developments and Fe2+ Uptake by Two Rice Varieties in an Iron Toxic Paddy Soil of Burkina Faso (West Africa). AS. 2016;07(11):783–804.

43. de Souza R, Meyer J, Schoenfeld R, da Costa PB, Passaglia LMP. Characterization of plant growth-promoting bacteria associated with rice cropped in iron-stressed soils. Ann Microbiol. 2015 Jun;65(2):951–64.

44. Chandwani S, Chavan SM, Paul D, Amaresan N. Bacterial inoculations mitigate different forms of iron (Fe) stress and enhance nutrient uptake in rice seedlings (Oryza sativa L.). Environmental Technology & Innovation. 2022 May;26:102326.

45. Sarhan MS, Hamza MA, Youssef HH, Patz S, Becker M, ElSawey H, et al. Culturomics of the plant prokaryotic microbiome and the dawn of plant-based culture media – A review. Journal of Advanced Research. 2019 Sep;19:15–27.

46. Armanhi JSL, de Souza RSC, Damasceno N de B, de Araújo LM, Imperial J, Arruda P. A Community-Based Culture Collection for Targeting Novel Plant Growth-Promoting Bacteria from the Sugarcane Microbiome. Front Plant Sci. 2018 Jan 4;8:2191.

47. Lewis WH, Tahon G, Geesink P, Sousa DZ, Ettema TJG. Innovations to culturing the uncultured microbial majority. Nat Rev Microbiol. 2021 Apr;19(4):225–40.

48. Riva V, Mapelli F, Bagnasco A, Mengoni A, Borin S. A Meta-Analysis Approach to Defining the Culturable Core of Plant Endophytic Bacterial Communities. Cann I, editor. Appl Environ Microbiol. 2022 Mar 22;88(6):e02537–21.

49. Sondo M, Wonni I, Klonowska A, Koïta K, Moulin L. Quantification of diversity sampling bias resulting from rice root bacterial isolation on popular and nitrogen-free culture media using 16S amplicon barcoding. Chang HX, editor. PLoS ONE. 2023 Apr 6;18(4):e0279049.

50. Mendez Del Villar P, Bauer JM. Rice in West Africa: Dynamics, policies and trends. Cahiers Agricultures. 2013 Sep;22(5):336–44.

51. Demont M. Reversing urban bias in African rice markets: A review of 19 National Rice Development Strategies. Global Food Security. 2013 Sep;2(3):172–81.

52. Mutiga SK, Rotich F, Were VM, Kimani JM, Mwongera DT, Mgonja E, et al. Integrated Strategies for Durable Rice Blast Resistance in Sub-Saharan Africa. Plant Disease. 2021 Oct 1;105(10):2749–70.

53. Barro M, Konate KA, Wonni I, Kassankogno AI, Sabot F, Albar L, et al. Assessment of Genetic Diversity of Rice in Registered Cultivars and Farmers’ Fields in Burkina Faso. Crops. 2021 Nov 3;1(3):129–40.

54. Wonni I, Ouedraogo L, Verdier V. First Report of Bacterial Leaf Streak Caused by *Xanthomonas oryzae* pv. *oryzicola* on Rice in Burkina Faso. Plant Disease. 2011 Jan;95(1):72–72.

55. Wonni I, Cottyn B, Detemmerman L, Dao S, Ouedraogo L, Sarra S, et al. Analysis of *Xanthomonas oryzae* pv. *oryzicola* Population in Mali and Burkina Faso Reveals a High Level of Genetic and Pathogenic Diversity. Phytopathology®. 2014 May;104(5):520– 31.

56. Barro M, Kassankogno AI, Wonni I, Sérémé D, Somda I, Kaboré HK, et al. Spatiotemporal Survey of Multiple Rice Diseases in Irrigated Areas Compared to Rainfed Lowlands in the Western Burkina Faso. Plant Disease. 2021 Dec 1;105(12):3889–99.

57. Odjo T, Diagne D, Adreit H, Milazzo J, Raveloson H, Andriantsimialona D, et al. Structure of African Populations of *Pyricularia oryzae* from Rice. Phytopathology®. 2021 Aug;111(8):1428–37.

58. Barro M, Wonni I, Simonin M, Kassankogno AI, Klonowska A, Moulin L, et al. The impact of the rice production system (irrigated *vs* lowland) on root-associated microbiome from farmer’s fields in western Burkina Faso. FEMS Microbiology Ecology. 2022 Aug 25;98(9):fiac085.

59. Barro M, Konate KA, Wonni I, Kassankogno AI, Sabot F, Albar L, et al. Assessment of Genetic Diversity of Rice in Registered Cultivars and Farmers’ Fields in Burkina Faso. Crops. 2021 Nov 3;1(3):129–40.

60. Barro M, Wonni I, Simonin M, Kassankogno AI, Klonowska A, Moulin L, et al. The impact of the rice production system (irrigated vs lowland) on root-associated microbiome from farmer’s fields in western Burkina Faso. FEMS Microbiology Ecology. 2022 Jul 22;fiac085.

61. Ranganayaki S, Mohan C. Effect of sodium molybdate on microbial fixation of nitrogen. Zeitschrift fur allgemeine Mikrobiologie. 1981;21(8):607–10.

62. Burbage DA, Sasser M. A medium selective for Pseudomonas cepacia. Phytophathology. 1982;72(6):706–706.

63. Döbereiner J, Day J. Associative symbiosis in tropical grasses: Characterization of microrganisms and dinitrogen fixing sites. In: Proc 1st Int Symp Nitrogen Fixation Washington. Newton WE, Nyman CJN (eds), Pullman, Washington State University Press; 1976. p. 518–38.

64. Baldani JI, Reis VM, Videira SS, Boddey LH, Baldani VLD. The art of isolating nitrogen-fixing bacteria from non-leguminous plants using N-free semi-solid media: a practical guide for microbiologists. Plant Soil. 2014 Nov;384(1–2):413–31.

65. Estrada-de los Santos P, Rojas-Rojas FU, Tapia-García EY, Vásquez-Murrieta MS, Hirsch AM. To split or not to split: an opinion on dividing the genus Burkholderia. Ann Microbiol. 2016 Sep;66(3):1303–14.

66. Wilson K. Preparation of Genomic DNA from Bacteria. In: Current Protocols in Molecular Biology. Ausubel, F.M., Brent, R., Kingston, R.E., Moore, D.D., Seidman, J.G., Smith, J.A. and Struhl, K., Eds. Wiley & Sons, New York; 1987.

67. Sarwar M, Kremer R. Determination of bacterially derived auxins using a microplate method. Letters in Applied Microbiology. 1995;20:282–5.

68. Gordon SA, Weber RP. COLORIMETRIC ESTIMATION OF INDOLEACETIC ACID. Plant Physiol. 1951 Jan 1;26(1):192–5.

69. Loper J, Schroth M. Influence of bacterial sources of indole-3-acetic acid on root elongation of sugar beet. Phytopathology. 1986;76:386–9.

70. Frey-Klett P, Chavatte M, Clausse M, Courrier S, Roux CL, Raaijmakers J, et al. Ectomycorrhizal symbiosis affects functional diversity of rhizosphere fluorescent pseudomonads. New Phytologist. 2005 Jan;165(1):317–28.

71. Lesmana M, Rockhill RC, Sutanti D, Sutomo A. An evaluation of alkaline peptone water for enrichment of Vibrio cholerae in feces. Southeast Asian J Trop Med Public Health. 1985 Jun;16(2):265–7.

72. Cappuccino JG, Sherman N. Microbiology: A Laboratory Manual. In: Microbiology: A Laboratory Manual (third ed). New York: Benjamin/Cummings Pub. Co.,; 1992. p. 125–79.

73. Nicholas K, Nicholas HJ. Genedoc: a tool for editing and annotating multiple sequence alignments. Distributed by the author at: http://www.psc.edu/biomed/ genedoc. 1997;

74. Saitou N, Nei M. The neighbor-joining method: a new method for reconstructing phylogenetic trees. Molecular Biology and Evolution [Internet]. 1987 Jul [cited 2023 Jan 30]; Available from: https://academic.oup.com/mbe/article/4/4/406/1029664/The-neighborjoining-method-a-new-method-for

75. Tamura K, Stecher G, Kumar S. MEGA11: Molecular Evolutionary Genetics Analysis Version 11. Battistuzzi FU, editor. Molecular Biology and Evolution. 2021 Jun 25;38(7):3022–7.

76. Letunic I, Bork P. Interactive Tree Of Life (iTOL) v5: an online tool for phylogenetic tree display and annotation. Nucleic Acids Research. 2021 Jul 2;49(W1):W293–6.

77. Cole JR, Wang Q, Fish JA, Chai B, McGarrell DM, Sun Y, et al. Ribosomal Database Project: data and tools for high throughput rRNA analysis. Nucl Acids Res. 2014 Jan;42(D1):D633–42.

78. Oksanen J, Simpson GL, Blanchet FG. vegan: Community Ecology Package. R package version 2.6-2 April 2022 [Internet]. The Comprehensive R Archive Network http://cran.r-project.org; 2022. Available from: http://cran.r-project.org

79. Liu H, Wang Z, Xu W, Zeng J, Li L, Li S, et al. *Bacillus pumilus* LZP02 Promotes Rice Root Growth by Improving Carbohydrate Metabolism and Phenylpropanoid Biosynthesis. MPMI. 2020 Oct;33(10):1222–31.

80. Duca D, Lorv J, Patten CL, Rose D, Glick BR. Indole-3-acetic acid in plant–microbe interactions. Antonie van Leeuwenhoek. 2014 Jul;106(1):85–125.

81. Li CY, Zhou YL, Ji J, Gu CT. Reclassification of Enterobacter oryziphilus and Enterobacter oryzendophyticus as Kosakonia oryziphila comb. nov. and Kosakonia oryzendophytica comb. nov. International Journal of Systematic and Evolutionary Microbiology. 2016 Aug 1;66(8):2780–3.

82. Singha KM, Singh B, Pandey P. Host specific endophytic microbiome diversity and associated functions in three varieties of scented black rice are dependent on growth stage. Sci Rep. 2021 Jun 10;11(1):12259.

83. Kang SM, Shahzad R, Khan MA, Hasnain Z, Lee KE, Park HS, et al. Ameliorative effect of indole-3-acetic acid- and siderophore-producing *Leclercia adecarboxylata* MO1 on cucumber plants under zinc stress. Journal of Plant Interactions. 2021 Jan 1;16(1):30–41.

84. Salomon MV, Purpora R, Bottini R, Piccoli P. Rhizosphere associated bacteria trigger accumulation of terpenes in leaves of Vitis vinifera L. cv. Malbec that protect cells against reactive oxygen species. Plant Physiology and Biochemistry. 2016 Sep;106:295– 304.

85. Jiang H, Feng Y, Zhao F, Lin X. Characteristics and Complete Genome Analysis of Bacillus asahii OM18, a Bacterium in Relation to Soil Fertility in Alkaline Soils Under Long-Term Organic Manure Amendment. Curr Microbiol. 2019 Dec;76(12):1512–9.

86. Feng Y, Chen R, Hu J, Zhao F, Wang J, Chu H, et al. Bacillus asahii comes to the fore in organic manure fertilized alkaline soils. Soil Biology and Biochemistry. 2015 Feb;81:186–94.

87. Ormeño-Orrillo E, Gomes DF, Del Cerro P, Vasconcelos ATR, Canchaya C, Almeida LGP, et al. Genome of Rhizobium leucaenae strains CFN 299T and CPAO 29.8: searching for genes related to a successful symbiotic performance under stressful conditions. BMC Genomics. 2016 Dec;17(1):534.

88. Divan Baldani VL, Baldani JI, Döbereiner J. Inoculation of rice plants with the endophytic diazotrophs Herbaspirillum seropedicae and Burkholderia spp. Biology and Fertility of Soils. 2000 Mar 3;30(5–6):485–91.

89. Coutinho BG, Passos da Silva D, Previato JO, Mendonça-Previato L, Venturi V. Draft Genome Sequence of the Rice Endophyte Burkholderia kururiensis M130. Genome Announc. 2013 May 2;1(2):e00225–12.

90. Timm CM, Campbell AG, Utturkar SM, Jun SR, Parales RE, Tan WA, et al. Metabolic functions of Pseudomonas fluorescens strains from Populus deltoides depend on rhizosphere or endosphere isolation compartment. Front Microbiol [Internet]. 2015 Oct 14 [cited 2022 Dec 21];6. Available from: http://journal.frontiersin.org/Article/10.3389/fmicb.2015.01118/abstract

91. Hol WHG, de Boer W, de Hollander M, Kuramae EE, Meisner A, van der Putten WH. Context dependency and saturating effects of loss of rare soil microbes on plant productivity. Front Plant Sci [Internet]. 2015 Jun 30 [cited 2023 Jan 25];6. Available from: http://journal.frontiersin.org/Article/10.3389/fpls.2015.00485/abstract

92. Compant S, Samad A, Faist H, Sessitsch A. A review on the plant microbiome: Ecology, functions, and emerging trends in microbial application. Journal of Advanced Research. 2019 Sep;19:29–37.

93. Becker M, Asch F. Iron toxicity in rice—conditions and management concepts. Z Pflanzenernähr Bodenk. 2005 Aug;168(4):558–73.

94. Otoidobiga CH, Keita A, Yacouba H, Traore AS, Dianou D. Dynamics and Activity of Iron-Reducing Bacterial Populations in a West African Rice Paddy Soil under Subsurface Drainage: Case Study of Kamboinse in Burkina Faso. AS. 2015;06(08):860– 9.

95. Chin KJ, Liesack W, Janssen PH. Opitutus terrae gen. nov., sp. nov., to accommodate novel strains of the division “Verrucomicrobia” isolated from rice paddy soil. International Journal of Systematic and Evolutionary Microbiology. 2001 Nov 1;51(6):1965–8.

96. Dianou D, Lopes J, Traoré AS, Lino AR, Moura I, Moura JG. Characterization of Desulfovibrio sp. isolated from some lowland paddy field soils of Burkina Faso. Soil Science and Plant Nutrition [Internet]. 2012 Jan 4; Available from: 10.1080/00380768.1998.10414468

97. Deng D, Zhang Y, Liu Y. A Geobacter strain isolated from rice paddy soil with higher bioelectricity generation capability in comparison to Geobacter sulfurreducens PCA. RSC Adv. 2015;5(55):43978–89.

98. Horino H, Ito M, Tonouchi A. Clostridium oryzae sp. nov., from soil of a Japanese rice field. International Journal of Systematic and Evolutionary Microbiology. 2015 Mar 1;65(Pt_3):943–51.

99. Zhao R, Farag IF, Jørgensen SL, Biddle JF. Occurrence, Diversity, and Genomes of “ *Candidatus* Patescibacteria” along the Early Diagenesis of Marine Sediments. Glass JB, editor. Appl Environ Microbiol. 2022 Dec 20;88(24):e01409–22.

100. Wang Y, Liu Z, Chen Q, Yi L, Xu Z, Cai M, et al. Isolation and characterization of novel Fusobacterium nucleatum bacteriophages. Front Microbiol. 2022 Nov 3;13:945315.

101. Farag IF, Youssef NH, Elshahed MS. Global Distribution Patterns and Pangenomic Diversity of the Candidate Phylum “Latescibacteria” (WS3). Löffler FE, editor. Appl Environ Microbiol. 2017 May 15;83(10):e00521–17.

102. Becraft ED, Woyke T, Jarett J, Ivanova N, Godoy-Vitorino F, Poulton N, et al. Rokubacteria: Genomic Giants among the Uncultured Bacterial Phyla. Front Microbiol. 2017 Nov 28;8:2264.

103. Sheremet A, Jones GM, Jarett J, Bowers RM, Bedard I, Culham C, et al. Ecological and genomic analyses of candidate phylum WPS −2 bacteria in an unvegetated soil. Environ Microbiol. 2020 Aug;22(8):3143–57.

104. Sait M, Hugenholtz P, Janssen PH. Cultivation of globally distributed soil bacteria from phylogenetic lineages previously only detected in cultivation-independent surveys: Cultivation of soil bacteria. Environmental Microbiology. 2002 Dec 9;4(11):654–66.

105. Janssen PH, Yates PS, Grinton BE, Taylor PM, Sait M. Improved Culturability of Soil Bacteria and Isolation in Pure Culture of Novel Members of the Divisions *Acidobacteria*, *Actinobacteria*, *Proteobacteria*, and *Verrucomicrobia*. Appl Environ Microbiol. 2002 May;68(5):2391–6.

106. Joseph SJ, Hugenholtz P, Sangwan P, Osborne CA, Janssen PH. Laboratory Cultivation of Widespread and PreviouslyUncultured SoilBacteria. Appl Environ Microbiol. 2003 Dec;69(12):7210–5.

107. Breznak JA, Canale-Parola E. Morphology and physiology of Spirochaeta aurantia strains isolated from aquatic habitats. Arch Microbiol. 1975;105(1):1–12.

108. Canale-Parola E. Physiology and evolution of spirochetes. Bacteriol Rev. 1977 Mar;41(1):181–204.

109. Oberhardt MA, Zarecki R, Gronow S, Lang E, Klenk HP, Gophna U, et al. Harnessing the landscape of microbial culture media to predict new organism–media pairings. Nat Commun. 2015 Oct 13;6(1):8493.

110. Liu S, Moon CD, Zheng N, Huws S, Zhao S, Wang J. Opportunities and challenges of using metagenomic data to bring uncultured microbes into cultivation. Microbiome. 2022 May 12;10(1):76.

111. Traoré A, Traoré K, Traoré O, Bado BV, Nacro BH, Sedogo MP. Caractérisation des systèmes de production à base de riz pluvial strict dans les exploitations agricoles de la zone Sud-soudanienne du Burkina Faso. Int J Bio Chem Sci. 2016 May 19;9(6):2685.

112. Sessitsch A, Pfaffenbichler N, Mitter B. Microbiome Applications from Lab to Field: Facing Complexity. Trends in Plant Science. 2019 Mar;24(3):194–8.

